# m^6^A-mediated Cell-cell Communication Controls Planarian Regeneration

**DOI:** 10.1101/2021.07.29.454253

**Authors:** Guanshen Cui, Jia-Yi Zhou, Xin-Yang Ge, Bao-Fa Sun, Ge-Ge Song, Xing Wang, Xiu-Zhi Wang, Rui Zhang, Hai-Lin Wang, Qing Jing, Yongliang Zhao, Magdalena J Koziol, An Zeng, Wei-Qi Zhang, Da-Li Han, Ying Yang, Yun-Gui Yang

**Author notes:** These authors contributed equally: Guanshen Cui, Jia-Yi Zhou, Xin-Yang Ge, Bao-Fa Sun, Ge-Ge Song, Xing Wang. Correspondence to (Y.-G.Y.); (Y.Y.); (D.-L.H.); (W.-Q.Z.).

## Abstract

Regeneration is the regrowth of damaged tissues or organs, a vital mechanism responding to damages from primitive organisms to higher mammals. Planarian possesses active whole-body regenerative capability owning to its vast reservoir of adult stem cells, neoblasts, thus provides an ideal model to delineate the underlying mechanisms for regeneration. *N*^6^-methyladenosine (m^6^A) regulates stem cell renewal and differentiation. However, how m^6^A controls regeneration at whole-organism level remains largely unknown. Here, we demonstrate that the depletion of m^6^A methyltransferase regulatory subunit *wtap* abolishes planarian regeneration, through regulating cell-cell communication and cell cycle. scRNA-Seq analysis unveils that the *wtap* knockdown induces a unique type of neural progenitor-like cells (NP-like cells), characterized by specific expression of the cell-cell communication ligand *grn*. Intriguingly, the depletion of m^6^A-modified transcripts *grn*/*cdk9* (or *cdk7*) axis rescues the defective regeneration of planarian without *wtap*. Overall, our study reveals an indispensable role of m^6^A-dependent cell-cell communication essential for whole-organism regeneration.

## Introduction

Regeneration is a process of replacing or restoring damaged or missing cells, tissues to full function, which widely exists in animal kingdoms^1^. For example, hydra can regenerate from tiny body fragments to entire organisms and zebrafish can regenerate large portions of heart^2^. For humans, some tissues also have regenerative potential throughout life, including liver and muscle^3^. Why such beneficial capability has gradually become restricted in higher organisms remains largely unknown, since the lack of regenerative ability can compromise the outcome of recovering from severe injury and disease^4^.

Planarians are being considered as a popular model system to study regeneration at whole-organism level^5,6^. The discovery of the planarian stem cell (neoblast) marker *piwi*^7^ and development of neoblast isolation methods^8^, has opened up opportunities to unravel the mechanisms of planarian regeneration. A number of pathways and regulatory factors have been identified to be essential for planarian regeneration, such as Wnt^9,10^ and EGFR signaling^11^. Especially along the advances of third generation DNA^12^ and single-cell RNA sequencing technologies^13,14^, accumulating evidences suggest that the neoblast is a source for regeneration^13–15^. In addition, some studies have revealed that epigenetic modifications also exert functions in planarian regeneration. Mihaylova et al. found that the COMPASS family MLL3/4 histone methyltransferases are essential for the differentiation and regeneration of the planarian^16^. In addition, the CREB-binding protein (CBP) and p300 family of histone acetyltransferases homologs *Smed*-CBP2 and *Smed*-CBP3 displayed distinct roles in stem cell maintenance and functions^17^.

*N*^6^-methyladenosine (m^6^A), the most abundant dynamic internal mRNA modification, has been shown to be an epi-transcriptomic marker playing diverse regulatory roles under physiological and/or pathological conditions^18^. Studies in various model systems have revealed that m^6^A regulates stem cell self-renewal and differentiation^19–22^. Depletion of either m^6^A methyltransferases or its demethylases dramatically affects gene expression profiles^23–26^. High-throughput sequencing technologies enable the detection of the m^6^A location within the transcriptome^23,27^. Subsequently, m^6^A has been identified in various RNA species, and has been implicated in stem cell biology, developmental and cancer biology^18^. For example, two separate studies showed that m^6^A regulates the transition of embryonic stem cells from a pluripotent state to a differentiated state. In this case, m^6^A selectively marks transcripts that code for key transcription factors involved in differentiation^19,28^. This demonstrates that m^6^A is a molecular switch that regulates stem cell differentiation, a fundamental mechanism in development and stem cell biology. Moreover, recent work showed that m^6^A regulates hematopoietic stem cell regeneration^29^ and axon regeneration in the mouse nervous system^30^. While those previous studies have provided important insights into the underlying molecular mechanism of m^6^A-mediated regulation of regeneration at cellular level, whether and how m^6^A is implicated in regeneration in tissues and entire organisms is unknown.

Here, we employed planarian *Schmidtea mediterranea* as the model to investigate the role of m^6^A in neoblast function and whole-body regeneration. The results illustrated that depletion of m^6^A methyltransferase regulatory subunit *wtap* abolished planarian regeneration, which is mainly mediated by *wtap*-associated m^6^A genes functioning in cell-cell communication and cell cycle through m^6^A-seq analysis. Further scRNA-Seq demonstrated a unique type of neural progenitor-like cells (NP-like cells) specifically expressing a cell-cell communication ligand *grn*. Intriguingly, depletion of *wtap* with any one of granulin (GRN), cyclin-dependent kinase 9 (CDK9) and CDK7, an signaling axis modified by m^6^A, rescues the defective regeneration of *wtap* knockdown planarian. Collectively, we reveal an indispensable role of m^6^A-dependent cell-cell communication for whole-organism regeneration. These findings improve our current understanding of the critical molecular events controlling regeneration, and have the potential to benefit future cell- or tissue-based replacement therapies.

## Results

### m^6^A methyltransferase complex upregulated during planarian regeneration

During planarian regeneration, principal component analysis (PCA) of all regeneration time points profiled shows that timepoints close to 6 hpa, 3 dpa, and 7 dpa occupy distinct positions during the regeneration cycle and display unique composition of Piwi-1 positive population and Piwi-1 negative population^31^, resembling the combination of the both processes of tissue regeneration from proliferating neoblasts (epimorphosis) and remodeling of the existing tissues (morphallaxis)^32^. In order to gain an overview of the gene expression dynamics from our own perspective during planarian regeneration, we performed RNA-seq at five different time points after amputation (0 hpa, 6 hpa, 3 dpa, 7 dpa, 11 dpa; hpa: hour post amputation; dpa: day post amputation) (Fig. 1a). We grouped all the expressed genes into 6 clusters according to their distinct expression pattern (Fig. 1b). Since m^6^A modification is catalyzed by three key components of the methyltransferases, including *mettl3*, *mettl14* and indispensable regulatory component *wtap*^33,34^, we found all three genes displayed up-regulated expression at 7 dpa and 11 dpa (Fig. 1c), and all belong to the cluster 5. Based on the Gene Ontology (GO) functional analysis, we found that genes in cluster 5 are enriched in signaling, cell communication and development related pathways, including growth factors and transcription factors such as *wnt2*, *egfr*, *sox2* and *pax6* (Fig. 1d). Further functional analysis reveals genes from this cluster are mostly involved in cell signaling and differentiation, as well as nervous system development (Fig. 1e). These results suggest a potential regulatory role of m^6^A in these pathways important for planarian regeneration.

**Figure 1.**
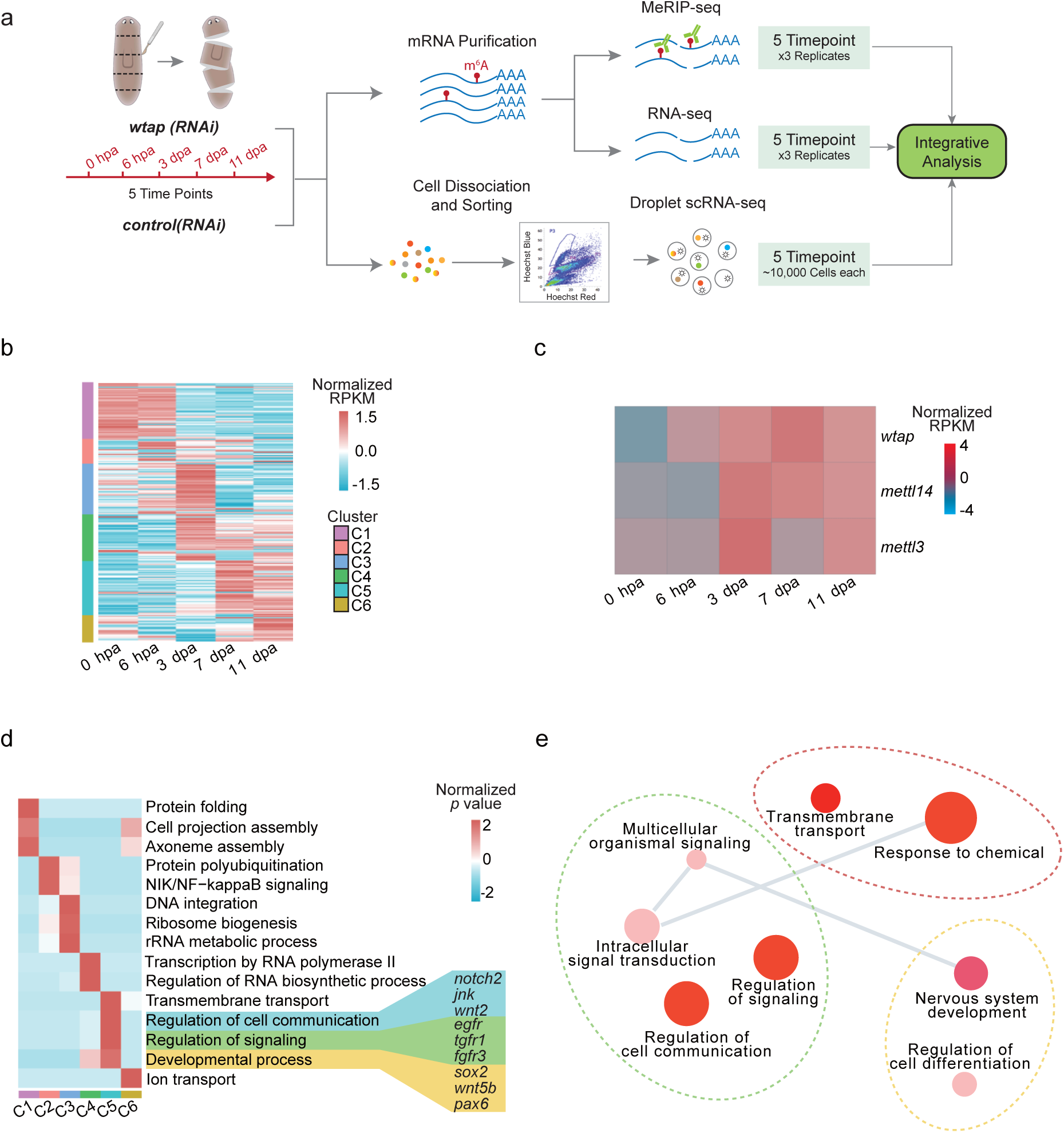
m^6^A methyltransferase complex upregulated during planarian regeneration. **a**, Experimental outline of multi-omics data including MeRIP-seq, bulk RNA-seq and single-cell RNA-seq. **b**, Heatmap showing the expression levels of genes within different expression pattern during 5 regeneration stages. **c**, Heatmap showing expression of the major components of m^6^A methyltransferase complex during planarian regeneration after amputation. **d**, Heatmap showing the enriched gene ontology for different clusters in Figure 1B. Color key represents z-score for -log_10_(*p* value). **e**, Gene ontology for genes of cluster 5 showing in **b.**

### m^6^A changes dynamically during planarian regeneration

To determine the regulatory role of m^6^A in planarian regeneration, we first tested the relative level of m^6^A in planarian mRNA, which shows a much higher level of over 0.5% relative to another modification on adenosine (*N*^1^-methyladenosine, m^1^A) detected using UHPLC-MRM-MS/MS (ultra-high-performance liquid chromatography-triple quadrupole mass spectrometry coupled with multiple-reaction monitoring) (Fig. 1a).

To obtain m^6^A landscape during planarian regeneration, we performed transcriptome-wide m^6^A mapping on samples of five time points (0 hpa, 6 hpa, 3 dpa, 7 dpa, 11 dpa) of regenerative process (Fig. 1a). We carried out MeRIP-seq to investigate the features and distribution dynamics of m^6^A in mRNA. In total, we identified 9,057-13,362 m^6^A peaks over all 5 different regeneration time points. We found that most genes have one m^6^A modification (Extended Data Fig. 1a). The total number of m^6^A peaks at each stage (especially at 6 hpa and 3 dpa) of regeneration is slightly higher than at 0 hpa (Extended Data Fig. 1b). Consistent with previous observations in mammals (Dominissini et al., 2012), we found that m^6^A peaks were markedly enriched near the stop codon. Also, the distribution pattern of m^6^A is similar among different regeneration time points (Fig. 2b). While all regeneration time points including the one before amputation (0 hpa) shares 6,385 genes that contains m^6^A peaks, there are 1,532 genes having their transcript being modified only after amputation (i.e. only in timepoints other than 0 hpa). Each time point also contains m^6^A modified genes unique to that particular time point during regeneration (Extended Data Fig. 1c, Supplementary Table 1).

**Figure 2.**
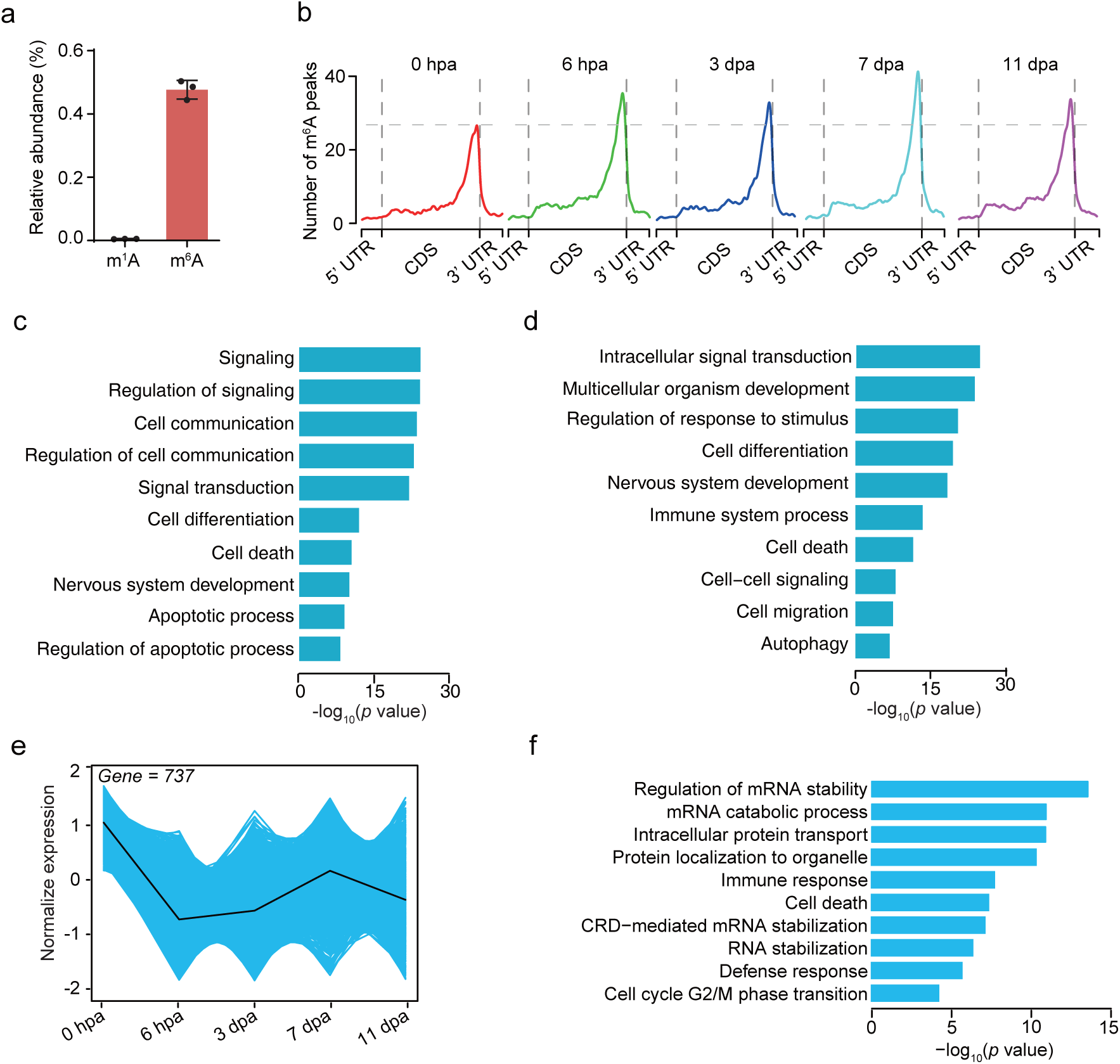
m^6^A changes dynamically during planarian regeneration. **a**, Barplot showing the abundance of m^1^A and m^6^A along mRNA. The methylation level of each RNA methylation was quantified *via* UHPLC-MRM-MS/MS. **b**, Metagene profiles of m^6^A peaks along transcripts with three non-overlapping segments (5’UTR, CDS, and 3’UTR) for 5 regeneration stages. **c**, Gene ontology terms for mRNAs with m^6^A modification which conserved at different regeneration stages showing in Figure S1C. **d**, Gene ontology enrichment for mRNAs with m^6^A modification which continuously up-regulated during regeneration. **e**, Line chart showing the fourth category of the mRNAs with m6A modification during regeneration, with gradual decreased expression from 0 hpa to 11 dpa. **e**, Gene ontology enrichment for genes shown in Figure 2E. See also **Extended Data Fig. 1.**

Gene Ontology enrichment analysis revealed that the 6,385 shared m^6^A-modified genes are associated with various essential biological processes, including regulation of signaling, regulation of cell communication and cell differentiation (Fig. 2c). This suggests that m^6^A may play important roles in these processes. All the m^6^A modified genes with increased expression during regeneration were related to signal transduction, cell differentiation, and nervous system development (Fig. 2d), further indicating an important role of m^6^A in these regeneration-related processes. The overlap of m^6^A modified transcripts and genes with different expression patterns indicate that the cluster 5 genes with upregulation at 7 dpa and 11 dpa have the highest overlap most with m^6^A (Extended Data Fig. 1d).

By classifying genes into up- or down- regulated genes during regeneration (Extended Data Fig. 1e-j, Fig. 2e-f), we found that m^6^A modified gene transcripts with up-regulated expression pattern similar to that of *wtap* were related to signal transduction, cell communication, cell differentiation, and nervous system development related pathways (Extended Data Fig. 1e-j, Fig. 2e-f). Conversely, down-regulated m^6^A genes during regeneration showed involvement in the regulation of mRNA stability and cell cycle G2/M phase transition related pathways (Fig. 2e-f), while the up-regulated m^6^A genes were related to regulation of signal transduction and system development. These results imply that m^6^A may participate in cell-cell communication and neuron development.

### m^6^A methyltransferase *wtap* depletion impairs regeneration in planarians

To further study the functional roles of m^6^A modification in planarian, we first performed the whole-mount *in situ* hybridization (WISH) of m^6^A methyltransferase regulatory subunit, *wtap*, and found that *wtap* specifically colocalizes with the neoblast marker *piwi*, especially at the posterior region, suggesting a regulatory role of m^6^A in neoblast (Fig. 3a-b). To study whether m^6^A modification is indispensable for the planarian regeneration, the key regulatory unit, *wtap*, was knocked down by RNA interference (RNAi), and then the worm was amputated at both anterior and posterior regions (Fig. 3c). Protein expression and mRNA levels of *wtap* were significantly reduced compared to the control, as shown by whole-mount immunostaining (Fig. 3d), Western blot (Extended Data Fig. 2a) and quantitative reverse-transcription PCR (qRT-PCR) (Extended Data Fig. 2b). Accordingly, the mRNA m^6^A level was reduced by nearly 80% at 7 dpa (Extended Data Fig. 2c).

**Figure 3.**
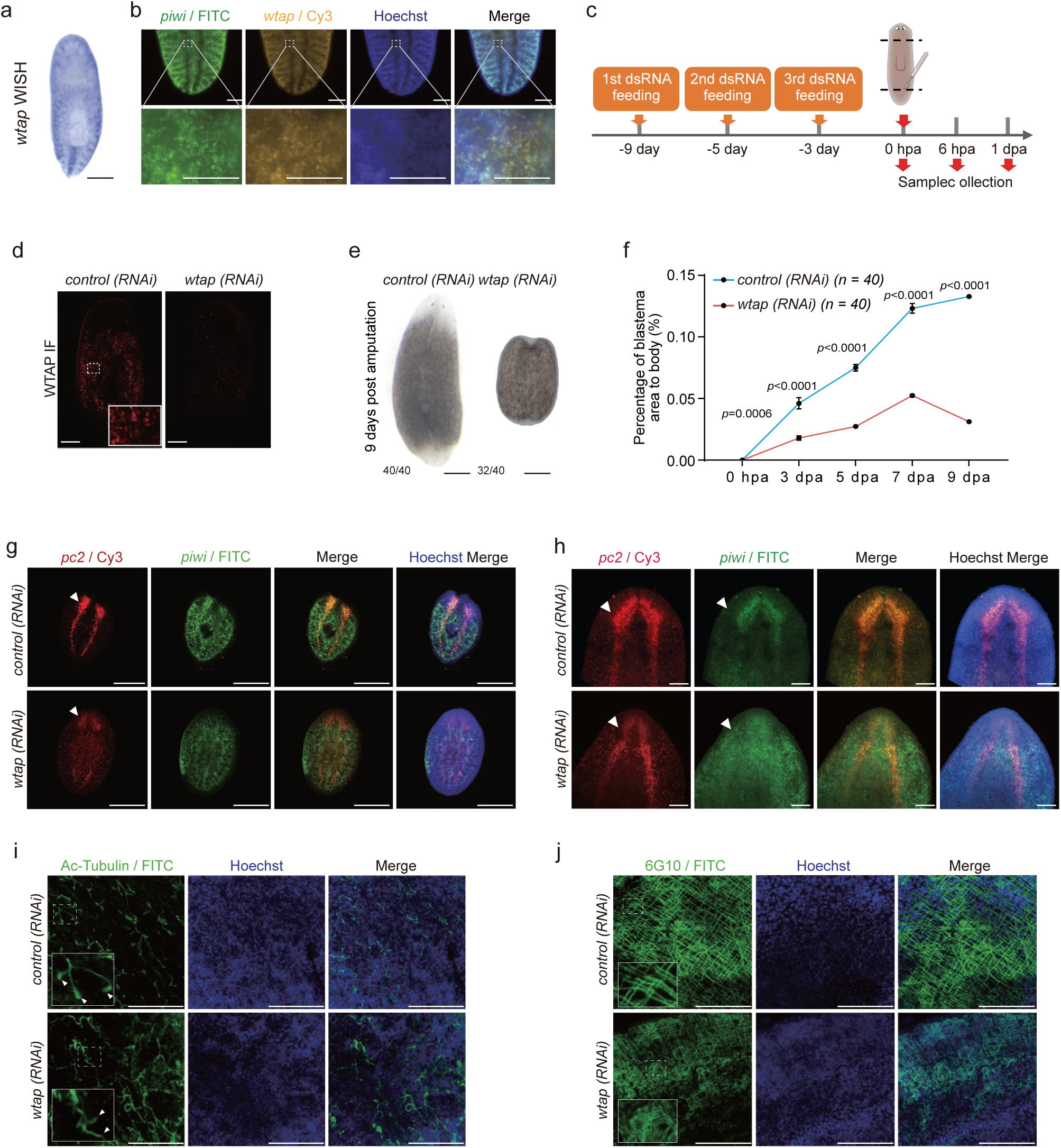
*wtap* depletion leads to regeneration defects. **a**, WISH showing the expression and localization of *wtap* transcripts in planarians. Scale bar, 300 µm. **b,** Whole-mount fluorescent *in situ* hybridization showing the expression and localization of *wtap* and *piwi* transcripts at posterior region. Scale bar, 200 µm (upper row), 100 µm (lower row). **c,** Schematic diagram showing the knockdown strategy and amputation position. **d**, Immunofluorescence (IF) showing expression and localization of WTAP proteins in control (*control*) and *wtap* depleted (*wtap* RNAi) planarians. (n ≥ 3) Scale bar, 200 μm. **e,** Bright-field image showing total body sizes for control (*control*) and *wtap* depleted (*wtap* RNAi) planarians at 9 dpa. Scale bar, 500 μm. Bottom left number, animals with phenotype of total tested. **f,** Percentage of blastema area to total body size in control (*control*) and *wtap* depleted (*wtap* RNAi) planarians. (n ≥ 3) Error bars represent standard deviation. Data were analyzed by the two-tailed unpaired Student *t-*test. ***, *P* < 0.001; ****, *P* < 0.0001. **g**, Whole-mount fluorescent *in situ* hybridization showing expressions and localizations of *pc2* (red) and *piwi* (green) along whole body of control (*control*) and *wtap* depleted (*wtap* RNAi) planarians at 5 dpa. Scale bar, 500 μm. **h,** Whole-mount fluorescent *in situ* hybridization showing expressions and localizations of *pc2* (red) and *piwi* (green) in head tissue of control (*control*) and *wtap* depleted (*wtap* RNAi) planarians at 5 dpa. Scale bar, 100 μm. **i,** Immunofluorescence showing the distribution of Ac-Tubulin protein in control (*control*) and *wtap* depleted (*wtap* RNAi) planarians at 7 dpa. (n ≥ 3) Scale bar, 200 μm. **j,** Immunofluorescence showing the distribution of 6G10 protein distribution in control (*control*) and *wtap* depleted (*wtap* RNAi) planarians at 7 dpa. (n ≥ 10) Scale bar, 200 μm. See also **Extended Data Fig. 2.**

Most importantly, *wtap* deficient planarians failed to fully regenerate the amputated tissue parts compared to the controls which regrew head and tail tissues at 9 dpa (Fig. 3e). Moreover, newly regenerated tissue in the *wtap* deficient planarian area was significantly smaller at 3 dpa than that of the control planarians (Extended Data Fig. 2d). The blastema formed at the wound region of *wtap* deficient planarians was also smaller and failed to grow the missing anterior and posterior tissues in comparison with the control planarians. (Fig. 3f). Especially at 7 dpa, when control planarians had regrown photoreceptors, the *wtap* knockdown planarians appeared to be still unable to give rise to any new tissues from its healed wound (Extended Data Fig. 3d). We noticed that *piwi* RNA expression did not change obviously in *wtap* knockdown planarians (Fig. 3g). Since *piwi* RNA is one of the most important and canonical marker of stem cells in planarians, *wtap* mediated m^6^A depletion may affect the neoblast function. At same time, the signatures of the level of regeneration are characterized by the regeneration of the nervous system containing a pair of ventral nerve cords (VNC) and the brain^35^. However, both parts were not regrown in *wtap* knockdown planarians, shown by whole-mount FISH staining of *pc2*, a neuropeptide processor expressed throughout the planarian central nervous system (Extended Data Fig. 2e), and in some cases the existing VNC disappeared (Fig. 3g-h, Extended Data Fig. 2e). Apart from failure to regenerate new tissues, the existing tissues were also abnormal upon *wtap* knockdown, such as the excretory system (Fig. 3i), and muscle fibers (Fig. 3j), indicated by the immunofluorescence staining of Ac-tubulin and 6G10, respectively.

Impaired regeneration and abnormal tissues morphology may occur due to the dysfunction of stem cells, neoblast or existing cell tissues. Thus, we investigated if proliferation and apoptosis is impaired in *wtap* knockdown planarians. We first performed BrdU pulse labeling assay and revealed no significant difference in global density of BrdU-positive cells between knockdown and control planarians at 5 dpa (Extended Data Fig. 2f, Extended Data Fig. 2g). But BrdU-labeled cells in *wtap* knockdown planarians were enriched near the amputation site, which was not found in control planarians, indicating the cell proliferation near the amputation site is more pronounced under *wtap* knockdown relative to control (Extended Data Fig. 2f). Then, we performed TUNEL assay to analyze apoptosis. The results showed a remarkable increase in cells undergoing apoptosis starting from 3 dpa in *wtap* deficient in comparison to control planarians (Extended Data Fig. 2h-i). Unchanged cell proliferation but increased cell death after *wtap* knockdown suggests that WTAP and m^6^A on cell cycle target transcripts regulate key components of the cell cycle. By doing so, this suggests a mechanism by which regeneration can be controlled.

### WTAP-mediated m^6^A controls cell cycle and cell-cell communication essential for regeneration

To investigate the underlying molecular mechanism for the defective planarian regeneration mediated by a decreased m^6^A level upon *wtap* knock-down, we performed m^6^A MeRIP-seq in control and *wtap* knockdown planarians (Fig. 1a). Even though the distribution of m^6^A along the CDS only slightly differs in *wtap* knockdown compared to controls, m^6^A peaks near stop codons become visibly reduced during the regeneration progression (Fig. 4a). We found that among the up-regulated genes belonging to cluster 5 (Fig. 1b), many genes showed a disordered expression pattern after *wtap* knockdown (Fig. 4b). Further GO functional analysis indicated that these genes are mainly related to regulation of signaling, cell communication and signal transduction associated pathways (Fig. 4b). We then analyzed the differentially expressed genes in each time point between control and *wtap* knockdown planarians. During the regeneration process, the number of both up-regulated and down-regulated genes went up starting from 0 hpa with highest number of genes changed at last stages, suggesting that the effect of *wtap* deficiency is stronger during the late stages than the early stages of regeneration process (Extended Data Fig. 3a). This indicates a sustaining, persisting and specific effect of *wtap* deficiency during the regeneration process. Gene set enrichment analyses (GSEA) revealed a marked up-regulation of the cell cycle signature and downregulation of cell-cell signaling in *wtap* deficient samples compared to control during regeneration (Fig 4c, Extended Data Fig. 3b). This suggests that *wtap* might be required for cell cycle and signaling transduction programs, as already observed before.

**Figure 4.**
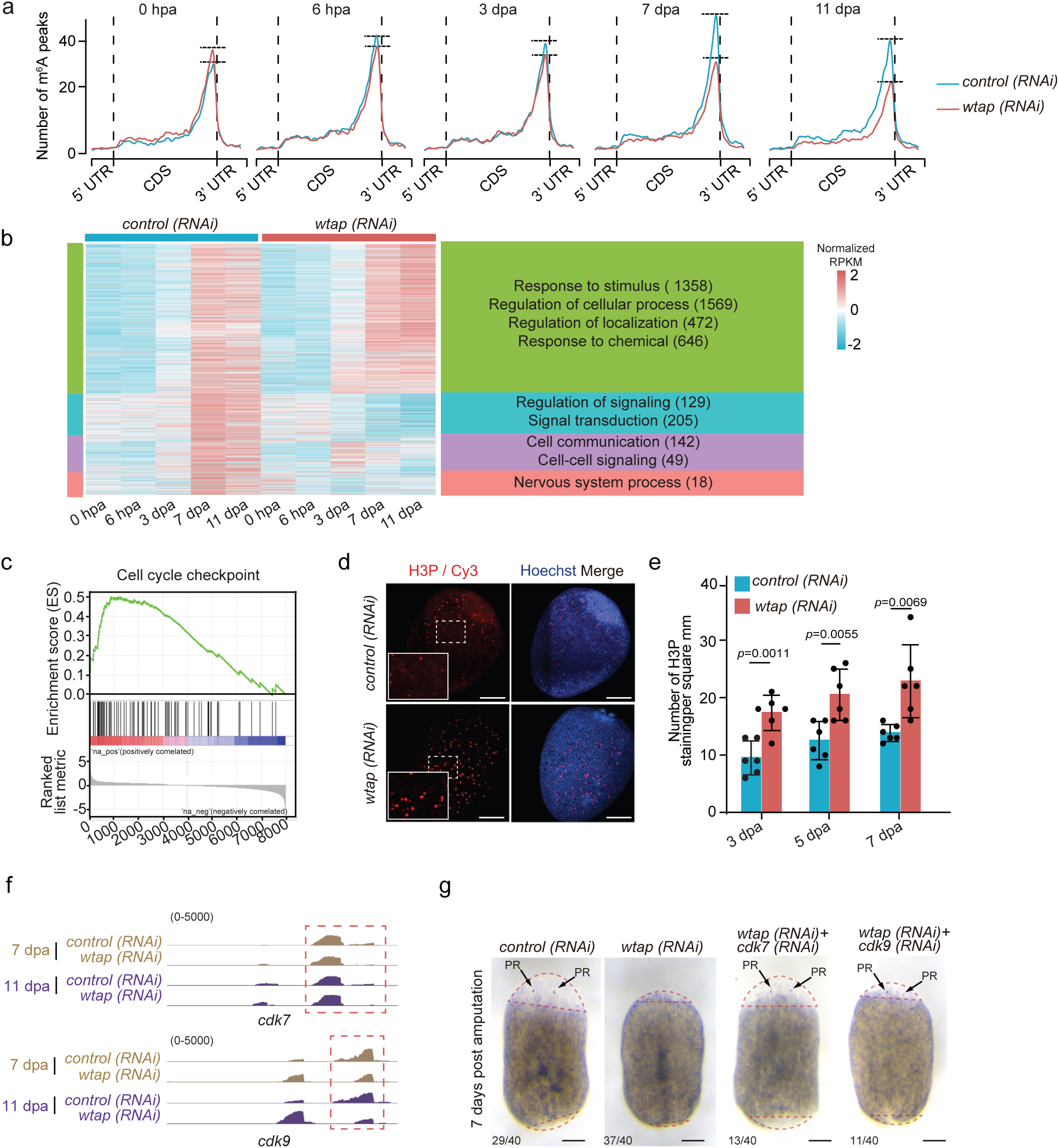
WTAP-mediated m^6^A controls cell cycle and cell-cell communication. **a**, Metagene profiles of m^6^A peaks along transcripts in both control (blue) and *wtap* knockdown (red) planarians for each regeneration stage after amputation (0 hpa, 6 hpa, 3 dpa, 7 dpa, 11 dpa). **b**, For genes from cluster 5 in Figure 1B, the expression level during 5 regeneration stages after amputation were displayed for both *control* and *wtap* knockdown planarians. Then those genes were separated into four new sub-groups based on expression features and the enriched GO terms were shown for each sub-group. **c**, GSEA plots evaluating the changes in cell cycle checkpoint pathway upon *wtap* depletion. Normalized *P* value < 0.01. **d**, Immunofluorescence showing the distribution of H3P protein in normal (*control*) and WTAP depleted (*wtap* RNAi) planarians at 5 dpa. Scale bar, 200 μm. **e**, Statistical analysis for H3P immunostaining. Error bars represent standard deviation. Data were analyzed by the two-tailed unpaired Student *t-*test. **, *P* < 0.01. **f,** Integrative Genomics Viewer (IGV) tracks displaying the distributions of reads from MeRIP-seq (red) and RNA-seq (blue) along *cdk7 and cdk9* in both *control* (top) and *wtap* knockdown (bottom) planarians at 3 dpa, 7 dpa and 11 dpa. **g**, Bright-field images showing the phenotypes of *control*, *wtap*, *cdk7*+*wtap* and *cdk9*+*wtap* knockdown planarians at 7 dpa. Scale bar, 100 μm. Bottom left number, number of animals with phenotype versus total number tested. See also **Extended Data Fig. 3.**

To further validate the effect of *wtap* knockdown on cell cycle, we performed immunofluorescence analysis using phosphorylated histone 3 (H3P), a typical marker for G2/M phase, and found a significant increase in the percentage of G2/M phase cells upon *wtap* knockdown (Fig 4d-e). One of key regulator of G2/M phase, Cyclin B1, was shown to have elevated level in whole-mount FISH staining during regeneration (Extended Data Fig. 3e). Therefore, combined with the result that no change in overall proliferation was observed but an increase in apoptosis upon *wtap* knockdown, the results suggest that *wtap* deficiency drives cells into the cell cycle resulting in G2/M phase stalling and eventually cell death. We further observed that the mRNA expression of *Cyclin-dependent kinase 7* (*cdk7*) and *cyclin-dependent kinase 9* (*cdk9*) was up-regulated upon *wtap* knockdown, while the level of m^6^A modification decreased (Fig. 4f). Genes related to cell-cell communication, such as *notum* and *kcnd2*, were down-regulated and their m^6^A level also decreased after *wtap* knockdown (Extended Data Fig. 3c-d). Furthermore, double knockdown of *cdk7* or *cdk9* with *wtap* rescues the *wtap* knockdown phenotype (Fig 4f), while neither of *cdk7* nor *cdk9* knockdown affects regeneratio (Extended Data Fig. 3f). This indicates that *wtap* knockdown phenotype is mediated by *cdk7* or *cdk9*. Taken together, the results show that WTAP controls planarian regeneration through m^6^A-modified genes functioning in cell-cell communication and cell cycle.

### scRNA-seq unveils impact of WTAP on specific cell types essential for regeneration

To decipher the detailed mechanism of m^6^A in regulating planarian regeneration, we conducted single-cell transcriptome sequencing of planarian *Schmidtea mediterranea* from five time points (0 hpa, 6 hpa, 3 dpa, 7 dpa, 11 dpa), in control and *wtap* knockdown samples. In total, 86,758 cells were successfully detected from all sequencing datasets of all samples. These were pooled together to generate a whole cell-type specific atlas. To identify individual cell types, we first performed unsupervised clustering using highly variable genes, and obtained 116 clusters by the t-distributed stochastic neighbor embedding method (t-SNE). We then elucidated the cell type identity of each cluster by identifying marker genes and compared them to markers in previous studies (Extended Data Fig. 4a). Overall, seven cell type groups such as neoblast, neuron, muscle, gut, secretory, epidermal and parenchymal cells were identified (Fig. 5a, Supplementary Table 2).

**Figure 5.**
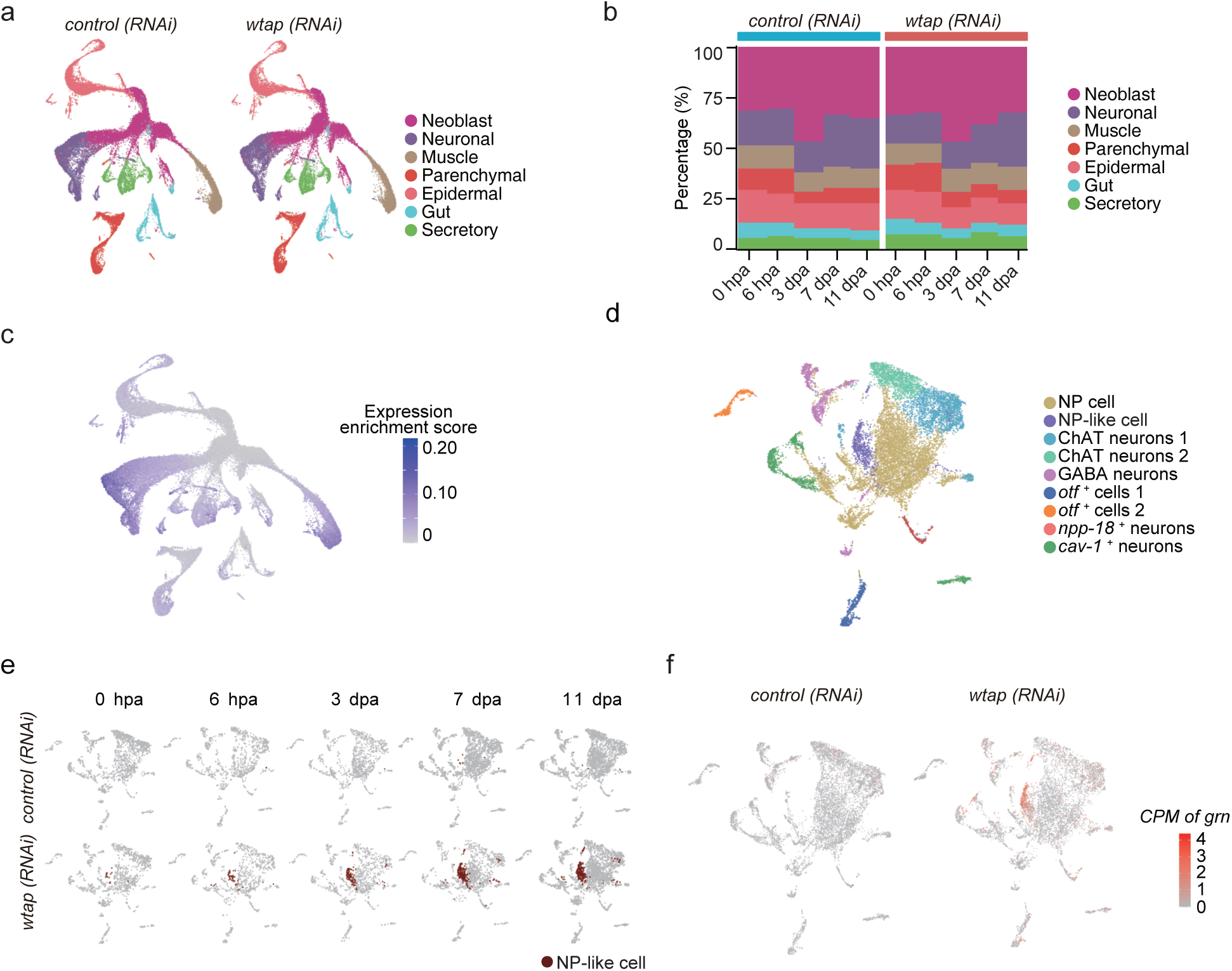
Single-cell atlas unveils cell-type specific regulation of WTAP essential for regeneration. **a**, Based on single-cell RNA-seq data, Uniform Manifold Approximation and Projection (UMAP) detect 7 different cell types in both control (left) and *wtap* knockdown (right) planarians during all 5 regeneration stages. **b**, Percentage of cells allocated to different cell types over time. **c**, Expression score of transcripts with m^6^A modifications shown in the cluster 5 of Figure 1B. **d**, UMAP analysis detects 9 sub-clusters of neuronal cells identified in **a. e**, UMAP plot of control versus *wtap* knockdown following amputation identifies NP-like cells in *wtap* knockdown planarians but that are absent in controls. **f**, Expression level of *grn* in neuronal cells. Normal (*control*) planarian on the left, *wtap* depleted (*wtap* RNAi) on the right. See also **Extended Data Fig. 4.**

In order to explore the regulation of m^6^A in cell lineage formation during planarian regeneration, we compared the cell-cell composition within control and *wtap* knockdown planarians. We found that the proportion of neural cells changes during regeneration. At 7 dpa, the proportion of regenerated neuronal cells in control planarians is about 25.0%, while it was 18.7% in *wtap* knockdown planarians. Notably, about one third of neuronal cells could not repopulate normally in *wtap* knockdown planarians (Fig. 5b, Supplementary Table 3). Except neuronal cells, the distribution of the other cell types between control and *wtap* knockdown samples was similar. In order to explore the cellular functional changes after *wtap* knockdown, we compared the control and *wtap* deficient samples of each cell type and found that epidermal and neoblast contain more up-regulated genes than the other five cell types. GO analysis revealed that during the regeneration, after *wtap* knockdown, the up-regulated genes in epidermal were enriched in nucleic acid metabolic process and cell cycle related functions (Extended Data Fig. 4b), and mainly related to cell cycle regulation functions in neoblast (Extended Data Fig. 4c). In addition, the down-regulated genes are mainly distributed in epidermal, neoblast and neuronal cells. Moreover, the down-regulated genes in epidermal cells are predominantly involved in many essential biological processes, including establishment of tissue polarity, translation, and ATP biosynthetic process. (Extended Data Fig. 4d). The function of down-regulated genes in neoblast is mainly associated with RNA splicing, DNA replication initiation, protein folding, etc. (Extended Data Fig. 4e). Neuronal cells are mainly associated with protein folding, neurotransmitter secretion and signal release from synapse functions (Extended Data Fig. 4f). These data suggest that *wtap* knockdown affects cell cycle and signal transduction pathways of neoblast and neuronal cells, which are important for planarian regeneration.

### WTAP-mediated m^6^A depletion increases GRN levels, which controls cell-cell communication essential for planarian regeneration

The results of single-cell data showed that the proportion of neural system showed dominant changes after *wtap* knockdown, and the m^6^A modified genes with the same expression trend as the known m^6^A methyltransferases were enriched in the neuronal cells (Fig. 5c). In addition, planarians could not regenerate nerve cord structure after *wtap* knockdown (Fig. 3Gg-h). In order to explore the underlying mechanisms for abnormal regeneration of the planarian nervous system, we further classified neuronal cells according to the reported marker ^14^. We clustered neuronal cells into 9 subtypes, neural progenitor cells (NPCs), NP-like cells, ChAT neurons 1, ChAT neurons 2, GABA neurons, *otf* ^+^ cells 1, *otf* ^+^ cells 2, *npp-18*^+^ neurons, *cav-1*^+^ neurons (Figure 5D). Among them, we identified a unique NP-like cell cluster during regeneration after *wtap* knockdown, which gradually increases during regeneration but is undetectable in the control planarians (Fig. 5e). We further identified *grn* and *ubiqp* as specific markers of NP-like cells (Fig. 5f Extended Data Fig. 5a, Supplementary Table 4).

To further investigate the functions of NP-like cells in planarian regeneration, we analyzed the effect of *wtap* knockdown on *grn* expression. First, the IGV and enrichment analyses showed that the level of m^6^A modification on *grn* transcript was decreased (Fig. 6a and Extended Data Fig. 5b). Consistently, WTAP-mediated m^6^A depletion leads to a drastic increase in *grn* and *ubiqp* expression in *wtap* deficient planarians (Extended Data Fig. 5c-d). Furthermore, through both whole-mount fluorescent *in situ* hybridization and immunofluorescence assays, we observed an increase in *grn* mRNA and protein levels (Fig. 6b-c). It has been reported that Progranulin (GRN) is a secretory protein important for neuronal functions^36–38^ and can exert function over a distance^36,39^. Furthermore, GRN can bind to the positive transcription elongation factor b (P-TEFb) and inhibit the phosphorylation of carboxy-terminal domain (CTD) of RNA polymerase II, which leads to transcription repression^39^. Our single-cell sequencing data showed that NP-like cells communicate with other cell types through secreting GRN (Extended Data Fig. 5g). Moreover, *Cyclin T1 (ccnt1)*, *cdk7* and *cdk9* have relative higher transcript abundance in the neoblast than any other cell types in the *wtap* knockdown planarians (Extended Data Fig. 5h), suggesting that GRN excreted by NP-like cells likely acts on neoblast. We found that the double knockdown of *wtap* and *grn* remarkably rescued the regeneration failure of planarians induced by *wtap* knockdown (Fig. 6d and Extended Data Fig. 5e), even though *grn* knockdown did not affect regeneration (Extended Data Fig. 5f). At same time, other cell-cell communication related gene transcripts such as, *mtnr1b*, also seem to regulate planarian regeneration (Extended Data Fig. 5i). These data suggest that WTAP-mediated m^6^A controls GRN expression levels, which further influences the regeneration of planarians.

**Figure 6.**
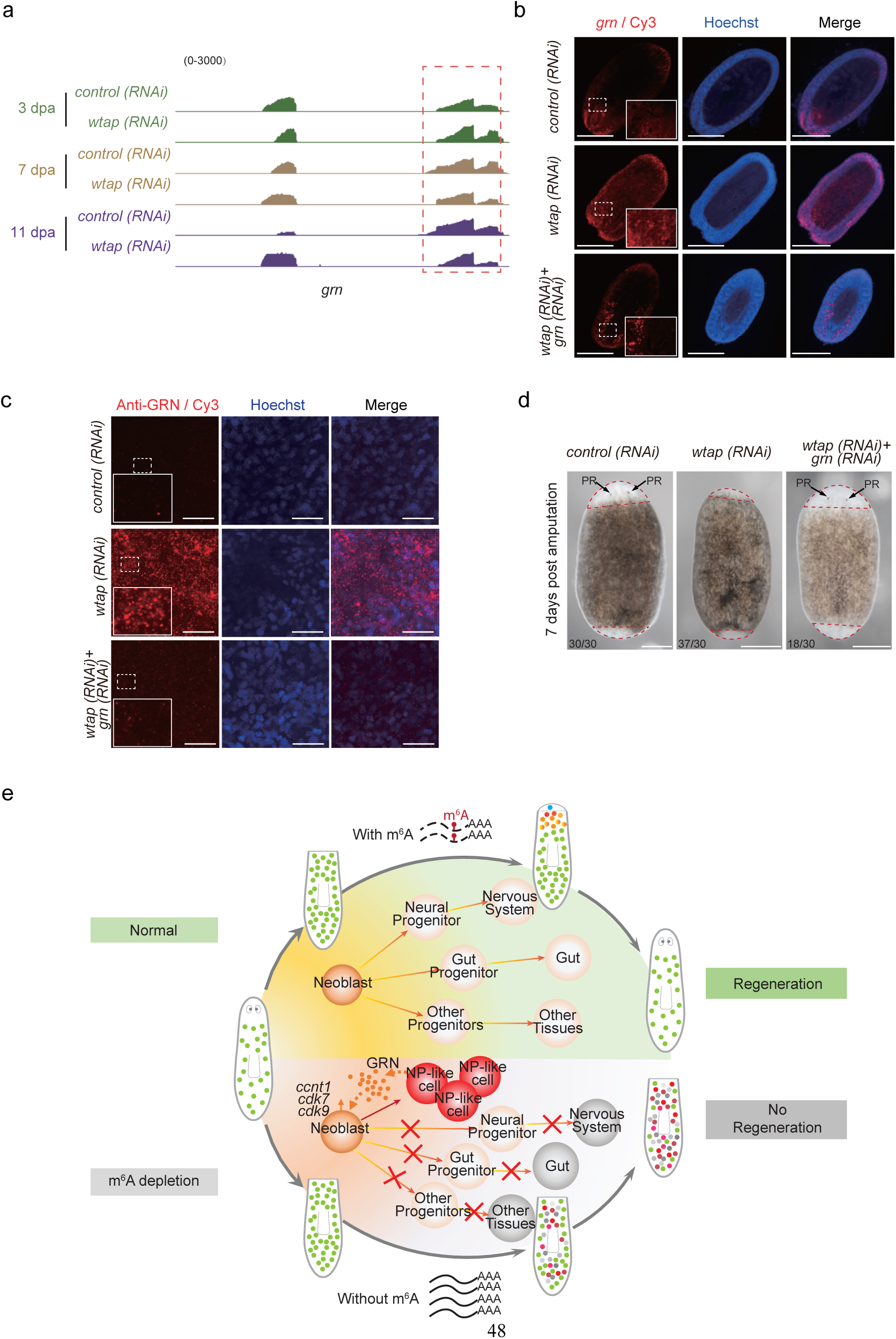
WTAP-mediated m^6^A depletion increases GRN levels, and influences cell-cell communication essential for planarian regeneration. **a**, IGV tracks displaying the distributions of reads from MeRIP-seq and RNA-seq along *grn* in both *control* (top) and *wtap* knockdown (bottom) planarians at 3 dpa, 7 dpa and 11 dpa. **b**, Whole-mount fluorescent *in situ* hybridization showing expressions of *grn* (red) in *control*, *wtap* and *grn*+*wtap* knockdown planarians at 7 dpa. Hoechst indicates DNA staining. Scale bar, 100 μm. **c**, Immunofluorescence showing the expression and localizations of GRN protein in *control*, *wtap* and *grn*+*wtap* knockdown planarians at 7 dpa. Hoechst indicates DNA staining. Scale bar, 200 μm. **d**, Immunofluorescence showing the expression and localizations of GRN protein in *control*, *wtap* and *grn*+*wtap* knockdown planarians at 7 dpa. Hoechst indicates DNA staining. Scale bar, 200 μm. **e**, Working model for WTAP-mediated m^6^A regulates planarian regeneration. See also **Extended Data Fig. 5.**

## Discussion

Regeneration is a dynamic and tightly-controlled process involving complex yet well-orchestrated gene regulation network. However, the potential significance of RNA m^6^A modification in regeneration is not well understood. In this study, we discovered for the first time that abundant RNA m^6^A modification exists in the planarian *Schmidtea mediterranea*, with a single high peak near stop codon region and enrichment at CDS and 3’ UTR region of mRNA which are similar as the features in most model organisms. Interestingly, this m^6^A peak increases along the regeneration process, and functional analysis further demonstrated that the m^6^A modified genes are related to pathways of cell-cell communications, nervous system development, mRNA stability and cell cycle G2/M phase transition, suggesting important regulatory roles of m^6^A modification in planarian regeneration. Moreover, m^6^A depletion by *wtap* knockdown led to a complete loss of regenerative ability of planarian attributing to the induction of cell cycle related genes *cdk7* and *cdk9* as well as the emergence of NP-Like cell cluster with aberrant expression of cell-cell communication related secretory gene-*grn,* identified by m^6^A sequencing analysis and rescue assays. Therefore, the signal axis of *wtap*-m^6^A-regulated *grn/cdk9/cdk7* determine the planarian regenerative potential through modulating cell-cell communication.

RNA m^6^A modification has been reported to regulate stem cell differentiation and tissue regeneration^19,21,28–30,40^. It is required for axon regeneration through promoting protein translation of regeneration-associated genes in the rodent model^30^. Meanwhile, deletion of m^6^A reader YTH domain-containing family protein 2 (YTHDF2) in mouse hematopoietic stem cells (HSC) was shown to facilitate HSC regeneration by enhancing stability of mRNAs related to both Wnt signal and survival^29^. In this study, using planarian *Schmidtea mediterranea* model system, we found the m^6^A peaks increase at CDS and 3’ UTR junction during the regeneration progression. Moreover, knockdown of m^6^A regulatory unit, *wtap*, resulted in a signification reduction in mRNA m^6^A level and abolished the regenerative capability of planarian to regrow missing head and tail. These findings suggest that *wtap*-mediated m^6^A is imperative for whole-organism regeneration of planarian.

Regeneration involves cell proliferation and the consequent generation of new tissues, which requires tight control of cell cycle^41^. In our study, we found cell cycle related genes such as *cdk7, cdk9* and *ccnt1* are mainly expressed in neoblasts and up-regulated upon *wtap* knockdown. CDK7 and CDK9 are members of the cyclin-dependent protein kinase family. CDK7 plays a double role as it activates several other cyclin dependent kinases in cell cycle and participates to the initiation of transcription through phosphorylating RNA polymerase II and other transcriptional machinery targets^42,43^. CDK9 was found to be an elongation factor for RNA polymerase II-directed transcription and can facilitate the transition from abortive to productive elongation with *ccnt1* which encodes a member of the highly conserved cyclin C subfamily^44^. In addition, the CDK9/CCNT1 complex is important in regulating specific differentiative pathways of several cell types, such as muscle cells, neurons and T cells^45^. The abnormal of cell cycle has been shown to be associated with the failure of planarian regeneration upon Hippo knockdown^46^. Previous studies showed that cell cycle gene CDK9 together with cyclin T1 serve as transcription elongation factor to facilitate transcription by phosphorylating the carboxy-terminal domain (CTD) of RNA polymerase II, and this process can be inhibited by GRN^39^; Moreover, *grn* inhibits the expression of *gata1* and further the differentiation of erythroid in zebrafish^47^. In support, we found induced expression of *grn, cdk7* and *cdk9* post *wtap* knockdown accompanied by impaired regeneration, suggesting that m^6^A modification are critical for planarian regeneration through destabilizing the transcripts of these three genes.

Importantly, accumulative studies revealed that regeneration is a complicated process involving cell-cell communication in various tissue types^48^. The flow of long-range patterning information in regenerative morphogenesis is crucial to control stem cell behavior *in vivo*^49^. For example, muscle regeneration in response to injury, is an non-specific inflammatory response to trauma, involving interaction between muscle and the immune system^48^. Specifically, muscle-myeloid cell communication is critical for the early, proliferative stage of muscle regeneration, where tumor necrosis factor (TNF) regulates differentiation of muscle progenitor cells (MPCs) *via* p38 mitogen-activated protein kinase pathway. Furthermore, growth factors, such as insulin-like growth factor I (IGF1), released by M1 macrophage stimulates growth of muscle progenitor cells at terminal differentiation stage of regeneration. During planarian regeneration, gap junctional communication (GJC) has been shown to regulate the construction of anterior-posterior A/P patterning, e.g. innexin-5 (Inx-5), inx-12 and inx-13 mediate the existence and properties of instructive, long-range signals from the central nervous system which controls pattern formation^49^. In our study, we identified a specific NP-like cell cluster secreting growth factor GRN after *wtap* knockdown, suggesting that GRN might selectively target the neoblast to inhibit their differentiation into neural progenitor cells and block the eventual formation of individual organs.

There are seven major cell types identified in planarian by scRNA-seq^14^. The most important cell type among them is the neoblast, which can provide the cellular source for robust planarian regeneration. This totipotent stem cell can differentiate into all other cell types after injury or stimulation. Soon after amputation, neoblast starts to proliferate and migrate to wound site to form regenerative blastemal tissue, which become the foundation for re-growing the missing tissue^6^. However, both blastema formation and remodeling of pre-existing tissue (morphallaxis) are required for the planarian regeneration^41^. These intricate processes require correct cell-cell communications and precise coordination between stem cells and pre-existing tissue to ensure the regeneration finished on the right track^6^. In our study, we identified a neural progenitor like cell cluster accumulated during the regeneration upon *wtap* knockdown, and further experiments demonstrated that the aberrant high expression of m^6^A modified gene, *grn*, secreted from this cell cluster disrupts the regeneration process. It is likely that such abnormal cell cluster is supposed to be a group of neural progenitor cells at special state during normal neural regeneration, which at same time is also critical for subsequent events of regeneration of other tissue types to proceed. Therefore, we speculate that this special state of neural progenitor cell cluster, as we amplified its cell number by *wtap* knockdown, triggers a critical checkpoint regulated during the entire regeneration process, that involves cell-cell communication mediated by GRN to ensure the correct path for regeneration, and *wtap* knockdown induced aberrant *grn* expression will stop the regeneration at this checkpoint.

As depicted in our model (Fig. 6e), during planarian homeostasis, the expression level of *grn* is held at moderate level to inhibit overgrowth, as equivalent to *grn* expression level before amputation at 0 hpa (Extended Data Fig. 5c). Upon injury, m^6^A modification selectively targets *grn* gene transcript for degradation, manifested as its reduced expression level after 6 hpa (Extended Data Fig. 5c), as well as transcripts of cell cycle related genes including *ccnt1*, *cdk7* and *cdk9*. m^6^A mediated down-regulation of these genes, especially the *grn*, dis-inhibits the polymerase II activity and overall transcription of the cell to promote the regeneration process, including the differentiation of neoblasts to the progenitor cells, and eventually to different tissue types. After *wtap* knockdown, *grn* transcript accumulates to form a unique cell cluster resulting in increased GRN secretion, which leads to the inhibition of neoblast differentiation and the failure of planarian regeneration.

Collectively, our study uncovered an essential role of WTAP-mediated m^6^A modification in regeneration of an entire organism. We further discovered that WTAP-mediated m^6^A modification is particularly important for the neural progenitor cell population. So far, this process has not been well understood. Our study may have uncovered a novel mechanism in regulating planarian regeneration that utilizes GRN to monitor and permit the regeneration happening correctly. Therefore, our study highlights the importance of cell-cell communication during the planarian regeneration, and provides important insights for future study in regenerative medicine.

## Methods

### Planarian culture

Animals Clonal asexual (CIW4) and sexual strains of *Schmidtea mediterranea* were maintained in Montjuïch salts as previously described (1.6 mM NaCl, 1.0 mM CaCl_2_, 1.0 mM MgSO_4_, 0.1 mM MgCl_2_, 0.1 mM KCl, and 1.2 mM NaHCO_3_ prepared in autoclaved Milli-Q water)^50^. Animals were fed weekly with homogenized pig liver. All animals, 3–6 mm in length, were starved 1 week before any experiments.

### Replication, size estimation, and randomization

For every assay, at least three independent replicates with a minimum of three animals per experiment were performed. For RNAi phenotype characterization, numbers of animals used are indicated in each panel. No sample size estimation was performed. Animals for all experiments were randomly selected. All animals were included in statistical analyses, and no exclusions were done. Images were randomized before quantifications.

### RNAi

Double stranded RNA (dsRNA) was synthetized by *in vitro* transcription as previously described. In summary, whole worm cDNA was generated and subcloned into a vector. *In vitro* transcription was performed using T7 polymerase (Promega, P2075). Planarians were fed by dsRNA mixed with pig liver paste three times every 3 days and were amputated into three fragments pre- and post-pharyngeal 24 hours after the last feeding. Primers used for the dsRNA synthesis are listed in Supplementary Table 5.

### BrdU labeling and whole-mount immunofluorescent staining

Animals were fed with 20 mg/mL BrdU (Sigma, 19160) as described before^7^. Specimens were fixed in 4% formaldehyde (FA) and antibody labelings were using rat anti-BrdU (Abcam, ab6326). Immunofluorescence staining was performed as described before^51^. Different treatments were used for different antibodies. Following antibodies were used: Phospho-histone H3 (Ser10) (H3P) (Abcam, ab32107). Muscle fibers were stained with the 6G10 antibody (1:1000, DSHB, 2C7; RRID: BDSC_9315). GRN protein was stained with anti-GRN (1:1000, Proteintech, 10053-1-AP). Planarians were killed in 5% N-acetyl cysteine (NAC) for 5 minutes at room temperature, and incubated in reduction buffer at 37°C for 8 minutes. After bleaching with 6% H_2_O_2_ overnight, planarians were washed with PBSTx (containing 0.3% Triton X-100) over 6 hours. The excretory system was stained with anti-acetylated-tubulin (1:1000, Sigma, T6793; RRID: AB_477585). Planarians were treated with 5% NAC for 5 minutes and 4% fixation for 20 minutes at room temperature followed by bleaching overnight and treated with proteinase K (1 µg/ml, Roche, 03115844001). After 2 hours of blocking with 1% BSA, planarians were incubated with primary antibody overnight, then washed with PBSTx (containing 0.3% TritonX-100) for more than 6 hours. Blocking with 1% BSA for 1 hour was performed before the fluorescent secondary antibody incubation overnight. After washing with PBSTx (containing 0.3% TritonX-100) for more than 6 hours, the slide was mounted with 80% glycerol containing Hoechst 33342 (10 µg/ml, Invitrogen, H3570).

### TUNEL assay

The TUNEL experiment was performed with ApopTag TUNEL Kit (Millipore, S7165) to determine apoptotic cell numbers in planarian following manufacturer’s instructions. 5% NAC (diluted in PBS) was used to remove the mucus coat of planarians. Planarians were fixed with 4% formaldehyde (with PBST containing 0.3% TritonX-100) for 30 minutes at room temperature. The samples were then bleached in 6% H_2_O_2_ (diluted in PBST) in direct light overnight. The animals were incubated with TdT reaction mixture for 4 hours at 37°C. The antibody was incubated for 4 hours at room temperature. Animals were washed every half an hour for 4 hours in PBST to reduce nonspecific labeling. Images were obtained with Leica SP8 confocal microscope. ImageJ was used for quantifications.

### Whole-mount *in situ* hybridization

DIG-labeled riboprobes were synthesized using an *in vitro* transcription kit (Roche, 11277073910), according to the manufacturer’s instructions. Animals were treated with 5% NAC for 5 min to clear epidermal mucus, and fixed in 4% formaldehyde in PBST for 30 min at room temperature. Then animals were dehydrated and stored in 100% methanol at -20°C for at least 1 h. The animals were bleached in 6% H_2_O_2_ under bright light overnight. Bleached samples were rehydrated through a graded series of methanol, and were incubated in probe mix at 56°C for 16 h. After the animals were washed and blocked, they were further incubated in anti-Digoxigenin-POD, Fab fragments (Roche, 11207733910; RRID: AB_514500) or anti-Digoxigenin-AP, Fab fragments (Roche, 11093274910; RRID: AB_514497) overnight at 4°C for 16 hours, followed by extensive washing. For Colorimetric whole-mount *in situ* hybridization, BCIP/NBT was used as substrates. For fluorescent development, a tyramide signal amplification system was used^52^. For double-color fluorescence *in situ* hybridization, POD inactivation was performed between signal developments in 100 mM NaN_3_ for 60 min. The primers used are listed in Supplementary Table 5.

### Total RNA extraction

Total RNA was extracted from planarians with 1 ml TRIzol^®^ reagent (Invitrogen, 15596018) through homogenizing on a tissue disruptor. mRNAs were purified from total RNAs using Dynabeads^®^ mRNA Purification Kit (Ambion, 61006) and subjected to TURBO™ DNase (Invitrogen) treatment at 37°C for 30 minutes and ethanol precipitation. After centrifugation and extensive washing with 75% ethanol, the mRNA was dissolved and quantified using Qubit 3.0 (Thermo Fisher).

### Western blot

Planarians were dissolved and homogenized in RIPA buffer (Cell Signaling Technology, 9806s) on a tissue disruptor. After incubation on ice for 10 minutes, the samples were centrifuged at 13,000 rpm for 30 minutes. Supernatant was collected and the Bradford method was used to measure protein concentrations. Western blot was performed as previously reported^53^ using the following antibodies: anti-WTAP antibody (1:500, Proteintech, 10200-1-AP; RRID: AB_2216349), anti-ß-actin antibody (1:1000, Cell Signaling Technology, 4967s; RRID: AB_330288). The density of each band was quantified with ImageJ (version 1.8.0).

### Quantitative reverse-transcription PCR (qRT-PCR)

To confirm knockdown efficiency and assess the relative RNA expression level, we conducted RT-qPCR. All RNA templates were digested with TURBO^TM^ DNase. cDNA synthesis was performed using the RevertAid^TM^ First Strand cDNA Synthesis Kit (Invitrogen, 18064014). Takara SYBR Premix Ex Taq (Takara) was used according to the manufacturer’s instructions and quantified by a CFX96 Real-Time PCR System (Bio-Rad). β-actin was used as an internal control. P-values were calculated using the two-tailed unpaired Student’s *t*-test. *, *P* <0.05; **, *P* <0.01; ***, *P* <0.001; ****, *P* <0.0001; ns, no significance. The primers used for qPCR are listed in Supplementary Table 5.

### UHPLC-MRM-MS/MS analysis

UHPLC-MRM-MS/MS analysis was performed as previously reported^54^. In brief, 200 ng mRNAs were purified from planarian total RNA at indicated time points. The mRNAs samples were digested overnight with 0.1 U Nuclease P1 (Sigma-Aldrich, N8630) and 1.0 U calf intestinal phosphatase (NEB, M0290) in a 0 µl reaction volume at 37°C. The mixture was subjected to UHPLC-MRM-MS/MS analysis to detect m^6^A after filtered the samples through ultra-filtration tubes (MW cutoff: 3 kDa, Pall, Port Washington, New York). The Agilent 1290 UHPLC system coupled with the 6495 triple quadrupole mass spectrometer (Agilent Technologies) was used for the analysis. UHPLC separation of mono-nucleosides was performed with a Zorbax Eclipse Plus C18 column (100 mm × 2.1 mm I.D., 1.8 µm particle size, Agilent Technologies). The mass spectrometer was operated in a positive ion mode. Three replicates were analyzed for each sample, with an injection volume of 5 µl. The amount of m^6^A and adenosine (A) was determined by using a calibration curve as a standard. Nitrogen was used for nebulization and desolvation, at 40 psi, with a flow-rate of 9 L/min, a source temperature of 300°C, capillary voltage 3,500 V, high purity nitrogen 99.999%. The ribonucleoside standards for m^6^A and A were purchased from TCI, China.

### RNA-seq and MeRIP-seq

mRNAs were purified from total RNAs using the Dynabeads^®^ mRNA purification kit (Ambion, 61006). cDNA libraries were made according to the TruSeq RNA Sample Prep Kit (Illumina, FC-122-1001) protocol. All samples were sequenced by Illumina Nova-seq with paired-end 150 bp read length. For m^6^A MeRIP-seq, the procedure is based on a published protocol^55^. Briefly, purified mRNA was fragmented to a size around of about 200 nt using the fragmentation reagent (Life Technologies, AM8740). 30 μl of protein A magnetic beads (Thermo Fisher Scientific, 10002D) was washed twice in 1 ml IP buffer (150 mM NaCl, 10 mM Tris-HCl [pH 7.5], 0.1% NP-40 in nuclease-free H_2_O), resuspended in 500 μl IP buffer mixed with 5 μg anti-m^6^A antibody (Millipore, ABE572) and incubated at 4 °C with gentle rotation for at least 6 hours. After 2 washes in IPP buffer (10 mM Tris-HCl [pH 7.4], 150 mM NaCl, 0.1% NP-40 in DEPC-treated), antibody-bead mixture was resuspended in 500 μl of the IPP reaction mixture containing 500ng fragmented mRNA, 100 μl of 5× IP buffer, and 5 μl of RNasin Plus RNase Inhibitor (Promega, N2611) and incubated for 2 hours at 4 °C. The beads were then washed twice each with 1 ml IPP buffer, low-salt IPP buffer (50 mM NaCl, 10 mM Tris-HCl [pH 7.5], 0.1% NP-40 in nuclease-free H_2_O) and high-salt buffer (500 mM NaCl, 10 mM Tris-HCl [pH 7.5], 0.1% NP-40 in nuclease-free H_2_O) for 10 min at 4°C. The beads were then eluted with 300 μl 0.5 mg/ml *N*^6^-Methyladenosine in IPP buffer (with RNasin) with gentle rotation at RT for 1 h. The m^6^A-modified RNAs were eluted using 200 μl of RLT buffer supplied in RNeasy Mini Kit (QIAGEN, 74106) for 2 minutes at room temperature. Supernatant was collected to a new tube while beads was pulled on magnetic rack. 400 μl of 100% ethanol was added to the supernatant. The mixture was applied to a RNeasy spin column and centrifuged at 13,000 rpm at 4°C for 30 seconds. The spin column was washed with 500 μl of RPE buffer, then 500 μl of 80% ethanol, and centrifuged at top speed for 5 minutes at 4°C to dry the column. m^6^A-modified RNAs were eluted with 10 μl nuclease-free H_2_O. For a second round of IP, eluted RNA was re-incubated with new protein A magnetic beads prepared with new anti-m^6^A antibody, followed by washes, elution and purification as above. Purified RNAs were used to construct library using the KAPA Standard RNA-Seq Kit according to the manufacturer’s instruction (KAPA, KR1139). Libraries were PCR amplified for 8-12 cycles and size-selected on the 8% TBE gel. Sequencing was carried out on the Illumina Nova 6000 platform according to the manufacturer’s instructions.

### Single-cell RNA-seq library construction

For single cell RNA-seq from 10x Chromium platform, Hoechst-stained and PI negative cells (200,000 cells) from wild-type animals, and from cells from *wtap* knockdown animals were collected on ice using Influx sorter. Approximately 9000 counted cells were loaded per channel. The libraries were made using the Chromium platform and Chromium Single Cell 3’ v3 chemistry. Sequencing libraries were loaded on an Illumina nova 6000 flowcell with two 150bp paired-end kits.

### RNA-seq data analysis

Adaptor sequences were trimmed off for all raw reads using the Cutadapt software (version 1.2.1)^56^. Reads that were less than 18 nt in length or contained an ambiguous nucleotide were discarded by Trimmomatic (version 0.30)^57^. The remaining reads were aligned to the *Schmidtea mediterranea* transcriptome smed_20140614 using Bowtie2 (v2.2.9)^58^ default parameters. Counts were calculated of the sum of reads of each transcript. R package ‘DEseq2’^59^ was used to identify differentially expressed transcripts. Genes with different expression pattern were defined by the K-means algorithm in MEV software^60^.

### MeRIP-seq data analysis

For MeRIP-seq, m^6^A-enriched peaks in each m^6^A immunoprecipitation sample were identified by MACS2 peak-calling software (version 2.0.10)^61^ with the corresponding input sample serving as control. MACS2 was run with default options except for ‘–nomodel’ to turn off fragment size estimation. A stringent cutoff threshold for a P value of 0.001 was used to obtain high-confidence peaks. In each stage, peaks with over 50% overlap in 3 biological replicates were used in the downstream analysis. The m^6^A level with different pattern were defined by the K-means algorithm in MEV software (version 4.4)^60^.

### Single cell RNA-seq data processing

Reads were processed using the Cell Ranger 3.0.0 pipeline^62^ with default and recommended parameters. FASTQ files generated from Illumina sequencing were aligned to the *Schmidtea mediterranea* transcriptome smed_20140614 using the STAR algorithm^63^. Next, Gene-Barcode matrices were generated for each individual sample by counting unique molecular identifiers (UMIs) and filtering non-cell associated barcodes. Only genes that can be translated into proteins were retained. We generated a gene-barcode matrix containing barcoded cells and gene expression counts. This output was then imported into the Seurat (v3.1.2)^64^ R toolkit for quality control and downstream analysis of single cell RNA-seq data. All functions were run with default parameters, unless specified otherwise. Low quality cells (<500 genes/cell, >6000 genes/cell, <15 cells/ gene and >20% mitochondrial genes) were excluded. In order to exclude multiple captures, which is a major concern in microdroplet-based experiments, DoubletFinder (version 2.0.2) (McGinnis et al., 2019) was employed to remove top N cells with the highest pANN score for each library separately, where N represents the doublet rates. Then all the datasets were merged using the “merge” function in Seurat.

### Identification of cell types and subtypes by nonlinear dimensional reduction (t-SNE)

The Seurat package implemented in R was applied to identify major cell types^64^. Highly variable genes were generated and used to perform PCA. Significant principle components were determined using JackStraw^64^ analysis and finally focusing on PCs 1 to 20. We grouped cell types based on their expression profile and matched them to known markers. To identify the subtype of neuronal cells, we used the reported markers of neuronal cells and also using “FindTransferAnchors” and “TransferData” function^64^ which provided by Seurat to predict the similarity between the subcluster and the reported cell types.

### Cell-cell communication analysis

In order to explore cell-cell communication networks via ligand–receptor interactions, we initially found the homologous genes of planarian and human genes by blastx (version 2.7.1)^65^. CellPhoneDB (v2.1.7)^66^ was used to identify the interaction among cells and all receptor-ligand pairs.

## Data availability

The datasets including RNA-seq, MeRIP-seq, and single cell sequencing data generated and analyzed during the current study are available in the Gene Expression Omnibus database under accession number GSE171253, and also the Genome Sequence Archive under accession number CRA004040 linked to the project PRJCA004712.

## Acknowledgements

This work was supported by grants from the Strategic Priority Research Program of the Chinese Academy of Sciences, China (XDA16010501), the National Natural Science Foundation of China (31625016, 31770872), and the National Key R&D Program of China (2018YFA0109700).

## Author contributions

Y.-G.Y. conceived this project, Y.-G.Y, D.-L.H., W.-Q.Z. and Y.Y. supervised the study and analyzed data; G.-S.C., X.-Y.G., G.-G.S., X.W. and X.-Z.W. performed the experiments; J.-Y.Z. and B.-F.S. performed bioinformatics analysis; R.Z. and H.-L.W. performed the UHPLC-MRM-MS/MS analysis; Y.-G.Y., Y.Y., D.-L.H., W.-Q.Z., G.-S.C., J.-Y.Z., X.-Y.G., G.-G.S., A.Z. and M.J.K. discussed and integrated the data, Q.J. provided the planarians, Y.-G.Y., Y.Y., G.-S.C., J.Y.Z., X.-Y.G., G.-G.S. and M.J.K. wrote the manuscript. All coauthors provided feedback on the final manuscript.

