## Supplementary Figures for "m^6^A-mediated Cell-cell Communication Controls Planarian Regeneration"

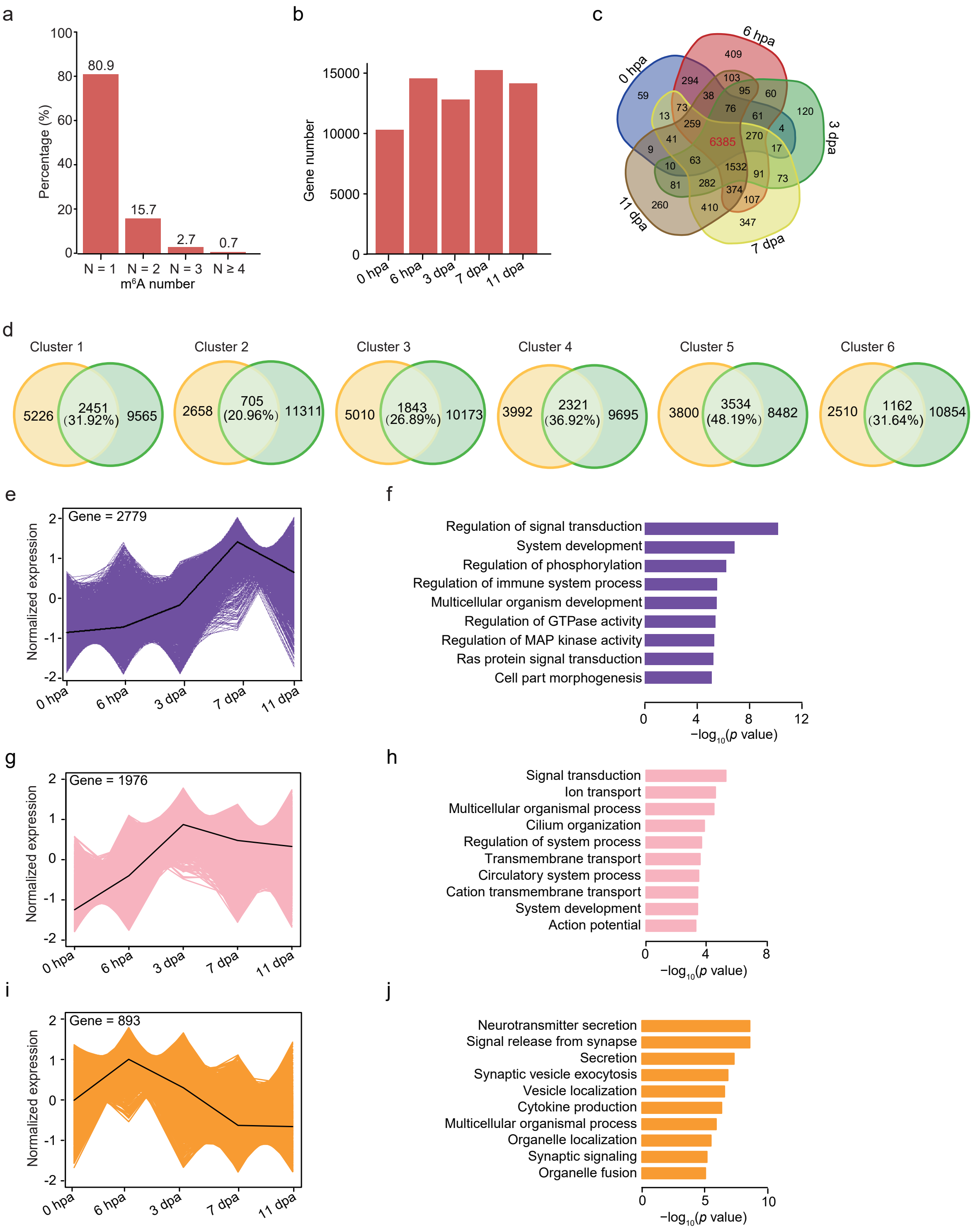
a
b
c
d
e
f
g
h
i
j
Percentage (%)
80.9
15.7
2.7
0.7
N = 1
N = 2
N = 3
N ≥ 4
m6A number
Gene number
0 hpa
6 hpa
3 dpa
7 dpa
11 dpa
6385
Cluster 1
5226
2451
(31.92%)
9565
Cluster 2
2658
705
(20.96%)
11311
Cluster 3
5010
1843
(26.89%)
10173
Cluster 4
3992
2321
(36.92%)
9695
Cluster 5
3800
3534
(48.19%)
8482
Cluster 6
2510
1162
(31.64%)
10854
Gene = 2779
Normalized expression
Regulation of signal transduction
System development
Regulation of phosphorylation
Regulation of immune system process
Multicellular organism development
Regulation of GTPase activity
Regulation of MAP kinase activity
Ras protein signal transduction
Cell part morphogenesis
−log10(p value)
Gene = 1976
Signal transduction
Ion transport
Multicellular organismal process
Cilium organization
Regulation of system process
Transmembrane transport
Circulatory system process
Cation transmembrane transport
System development
Action potential
Gene = 893
Neurotransmitter secretion
Signal release from synapse
Secretion
Synaptic vesicle exocytosis
Vesicle localization
Cytokine production
Multicellular organismal process
Organelle localization
Synaptic signaling
Organelle fusion

**Extended Data Fig. 1 | Changes of m$^6$A modified genes during regeneration, Related to Figure 2.**

**a**, Percentage of m$^6$A-methylated mRNAs with different peak numbers of m$^6$A peaks. **b**, The number of differentially expressed genes with m$^6$A modification in different periods. **c**, Overlap of m$^6$A-modified mRNAs in 5 regeneration stages. **d**, The overlap of mRNA from different clusters that shown in Figure 1B and total m$^6$A-modified mRNAs. **e**, Line chart showing the first category of the mRNAs with m$^6$A modification during regeneration, with increased expression from 0 hpa to 7 dpa and then decreased expression from 7 dpa to 11 dpa. **f**, Gene ontology enrichment for genes shown in Figure S2A. **g**, Line chart showing the second category of the mRNAs with m$^6$A modification during regeneration, with increased expression from 0 hpa to 3 dpa and then decreased expression from 3 dpa to 11 dpa. **h**, Gene ontology enrichment for genes shown in Figure S2C. **i**, Line chart showing the third category of the mRNAs with m$^6$A modification in different during regeneration, with increased expression from 0 hpa to 6 hpa and then decreased expression from 6 hpa to 11 dpa. **j**, Gene ontology enrichment for genes shown in Figure S2E.

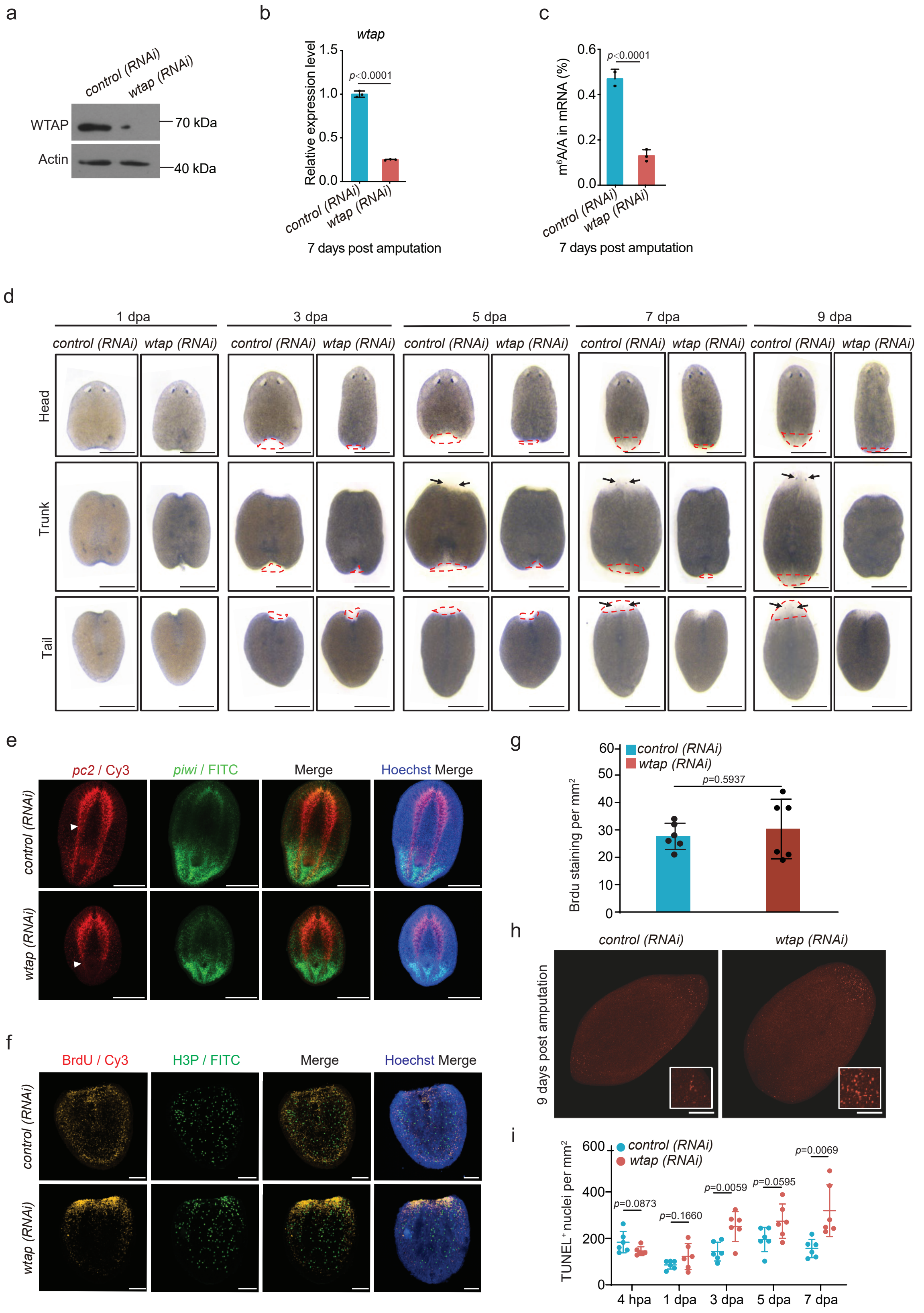

a
control (RNAi)
wtap (RNAi)
WTAP
70 kDa
Actin
40 kDa
b
wtap
Relative expression level
p<0.0001
7 days post amputation
c
m6A/A in mRNA (%)
p<0.0001
7 days post amputation
d
1 dpa
3 dpa
5 dpa
7 dpa
9 dpa
Head
Trunk
Tail
e
pc2 / Cy3
piwi / FITC
Merge
Hoechst Merge
f
BrdU / Cy3
H3P / FITC
Merge
Hoechst Merge
g
Brdu staining per mm2
p=0.5937
h
9 days post amputation
i
TUNEL+ nuclei per mm2
p=0.0873
p=0.1660
p=0.0059
p=0.0595
p=0.0069
4 hpa
1 dpa
3 dpa
5 dpa
7 dpa

**Extended Data Fig. 2 | Detection of *wtap* knockdown efficiency and $m^6A$ methylation levels, Related to Figure 3.**

**a**, Western blot showing the knockdown efficiency of WTAP. Actin is used as a loading control. **b**, Quantitative qPCR showing the relative expression of *wtap* upon *wtap* knockdown at 7 dpa. **c**, UHPLC-MRM-MS/MS showing $m^6A$ level of mRNA extracted from normal (*control*) and WTAP depleted (*wtap* RNAi) planarian at 7 dpa. Data were analyzed by the two-tailed unpaired Student *t*-test. ****, $p < 0.0001$; n.s., no significance. **d**, Bright-field images showing WTAP depleted (*wtap* RNAi) phenotype of head, trunk, tail tissue fragment at 1 dpa, 3 dpa, 5dpa, 7 dpa and 9 dpa. Scale bar, 500 µm. **e**, Whole-mount fluorescent *in-situ* hybridization showing the expression and distribution of *pc2* and *piwi* transcript of trunk tissue fragment at 5 dpa. Scale bar, 500 µm. **f**, Whole-mount fluorescent BrdU immunostaining of normal (*control*) and *wtap* depleted (*wtap* RNAi) planarian at 5 dpa. Scale bar, 300 µm. **g**, Quantification of $BrdU^+$ nuclei/$mm^2$ at 5 dpa. Error bars represent standard deviation. Data were analyzed by the two-tailed unpaired Student *t*-test. **h**, Whole-mount TUNEL assay showing the late apoptotic cells of normal (*control*) and *wtap* depleted (*wtap* RNAi) planarian 9 days post amputation. Scale bar, 200 µm. **i**, Quantification of $TUNEL^+$ nuclei/$mm^2$ at 4 hpa, 1 dpa, 3 dpa, 5 dpa, 7 dpa and 9 dpa. Error bars represent standard deviation. Data were analyzed by the two-tailed unpaired Student *t*-test. ***, $p < 0.001$; ns, no significance.

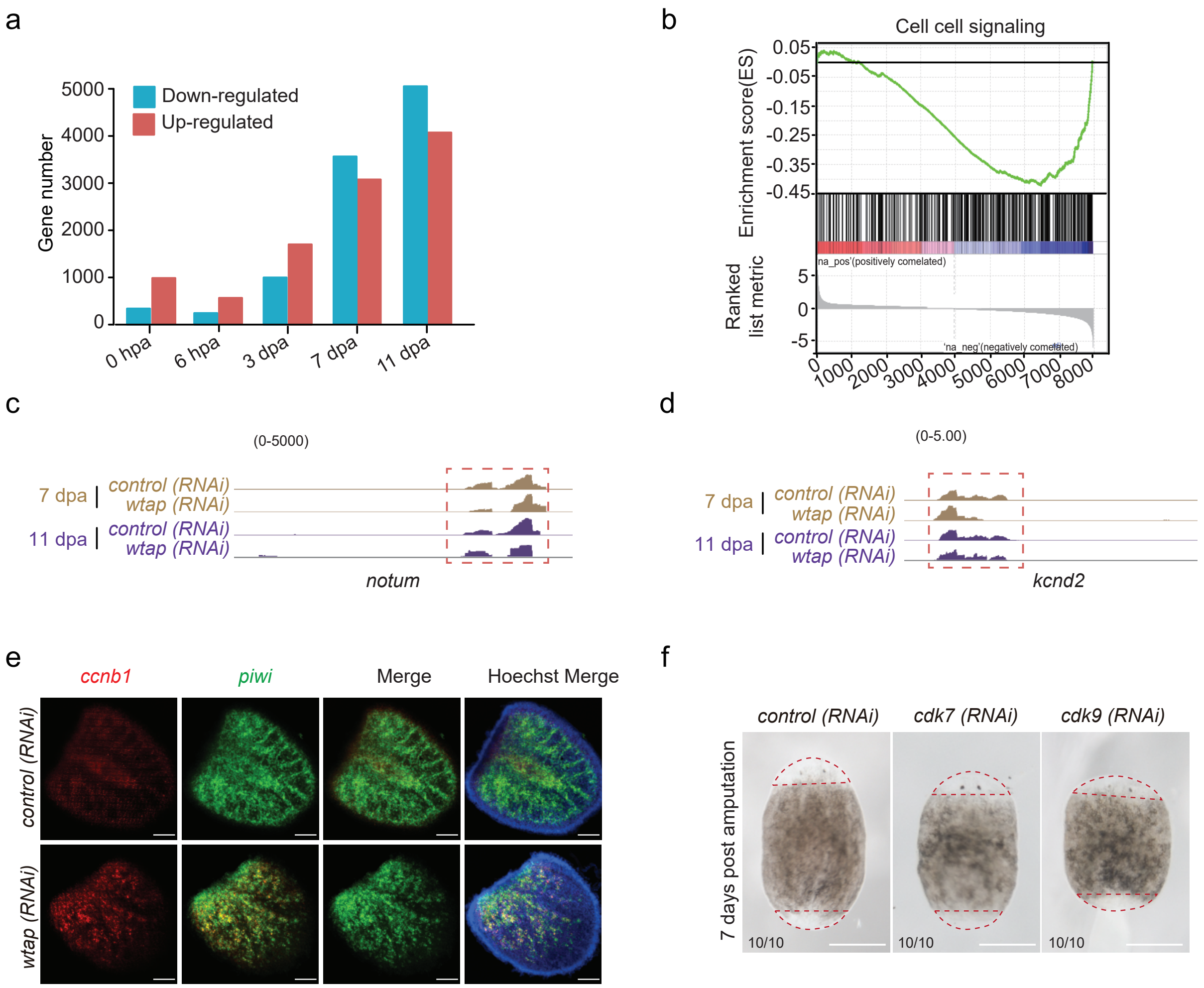
a
Down-regulated
Up-regulated
Gene number
0 hpa
6 hpa
3 dpa
7 dpa
11 dpa
b
Cell cell signaling
Enrichment score(ES)
Ranked list metric
'na_pos'(positively correlated)
'na_neg'(negatively correlated)
c
(0-5000)
7 dpa
11 dpa
control (RNAi)
wtap (RNAi)
notum
d
(0-5.00)
kcnd2
e
ccnb1
piwi
Merge
Hoechst Merge
control (RNAi)
wtap (RNAi)
f
control (RNAi)
cdk7 (RNAi)
cdk9 (RNAi)
7 days post amputation
10/10

**Extended Data Fig. 3 | WTAP-mediated m$^6$A controls cell cycle and cell-cell communication, Related to Figure 4.**

**a**, Bar plot showing the total number of up- and down- regulated transcripts that contain m$^6$A modifications under 5 timepoints following amputation. **b**, GSEA plots evaluating the changes in multicellular organismal signaling, compared WTAP depleted (*wtap* RNAi) with normal (*control*) planarian at 3 dpa. Normalized *P* value = 0.050. **c**, Integrative Genomics Viewer (IGV) tracks displaying MeRIP-seq and RNA-seq read distribution in *notum* mRNA of control and *wtap* depleted samples. **d**, IGV tracks displaying MeRIP-seq and RNA-seq read distribution in *kcnd2* mRNA of control and *wtap* depleted samples. **e**, Whole-mount fluorescent *in-situ* hybridization showing the expression and distribution of *ccndb1* and *piwi* transcript of trunk tissue fragment at 5 dpa. Scale bar, 500 µm. **f**, The phenotype of normal (control), *cdk7* depleted (*cdk7* RNAi) or *cdk9* depleted (*cdk9* RNAi) at 7 dpa. Bottom left number, animals with phenotype of total tested. Scale bar, 500 µm.

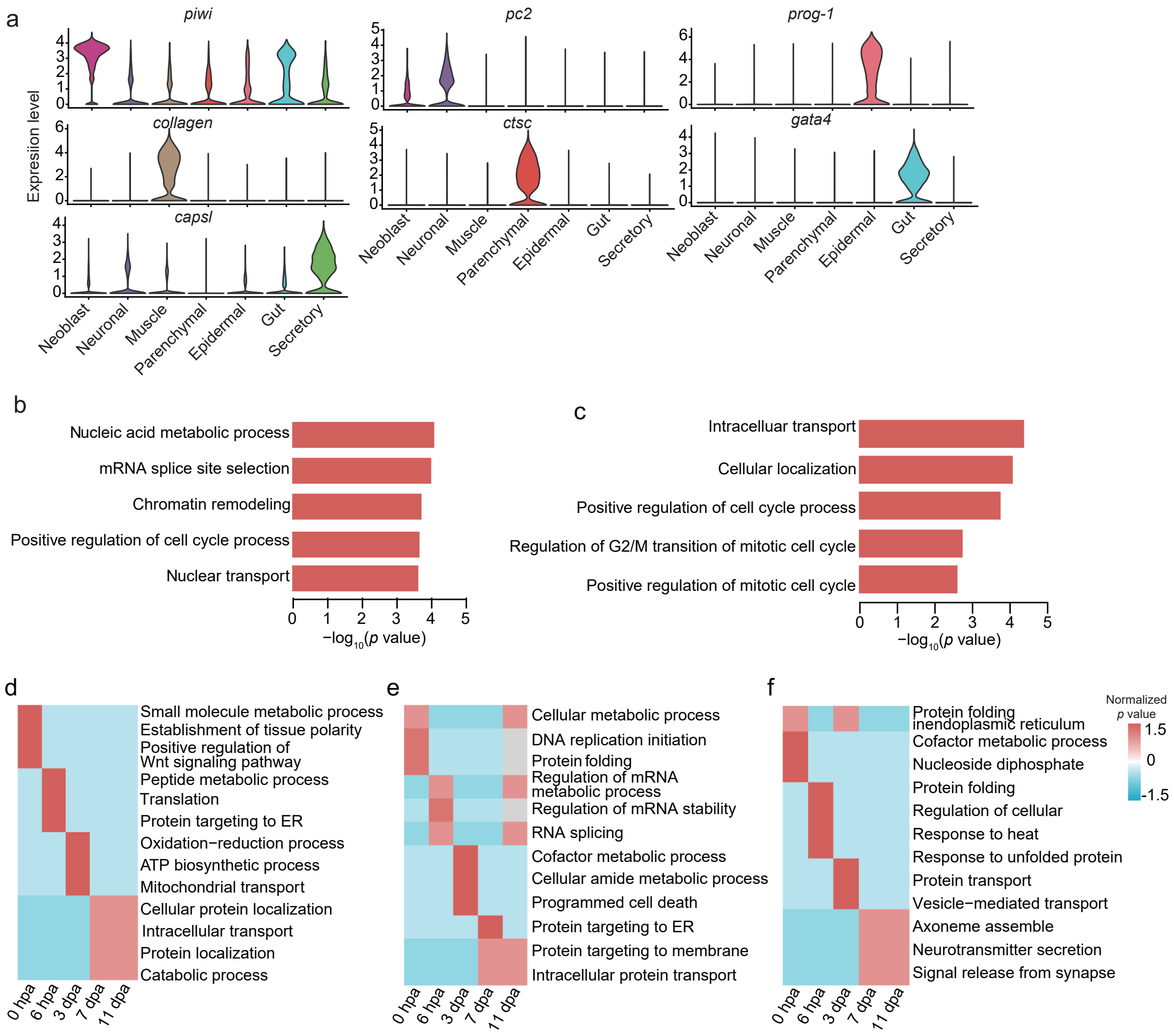
a
piwi
pc2
prog-1
collagen
ctsc
gata4
capsl
Expresion level
Neoblast
Neuronal
Muscle
Parenchymal
Epidermal
Gut
Secretory
b
Nucleic acid metabolic process
mRNA splice site selection
Chromatin remodeling
Positive regulation of cell cycle process
Nuclear transport
−log10(p value)
c
Intracelluar transport
Cellular localization
Positive regulation of cell cycle process
Regulation of G2/M transition of mitotic cell cycle
Positive regulation of mitotic cell cycle
−log10(p value)
d
Small molecule metabolic process
Establishment of tissue polarity
Positive regulation of Wnt signaling pathway
Peptide metabolic process
Translation
Protein targeting to ER
Oxidation−reduction process
ATP biosynthetic process
Mitochondrial transport
Cellular protein localization
Intracellular transport
Protein localization
Catabolic process
0 hpa
6 hpa
3 dpa
7 dpa
11 dpa
e
Cellular metabolic process
DNA replication initiation
Protein folding
Regulation of mRNA metabolic process
Regulation of mRNA stability
RNA splicing
Cofactor metabolic process
Cellular amide metabolic process
Programmed cell death
Protein targeting to ER
Protein targeting to membrane
Intracellular protein transport
f
Protein folding inendoplasmic reticulum
Cofactor metabolic process
Nucleoside diphosphate
Protein folding
Regulation of cellular
Response to heat
Response to unfolded protein
Protein transport
Vesicle−mediated transport
Axoneme assemble
Neurotransmitter secretion
Signal release from synapse
Normalized p value
1.5
0
-1.5

**Extended Data Fig. 4 | Functional analysis in different cell types upon *wtap* knockdown during regeneration, Related to Figure 5.**

**a**, Violin plots showing expression levels of specific markers for different cell types. **b**, Enriched gene ontology terms for up-regulated genes upon *wtap* knockdown at 3 dpa in neuronal cells. **c**, Enriched gene ontology terms for up-regulated genes upon *wtap* knockdown at 3 dpa in epidermal cells. **d**, Enriched gene ontology terms for down-regulated genes upon *wtap* knockdown in epidermal cells. **e**, Eenriched gene ontology terms for down-regulated genes upon *wtap* knockdown in neoblast cells. **f**, Enriched gene ontology terms for down-regulated genes upon *wtap* knockdown in neuronal cells at different regeneration stages.

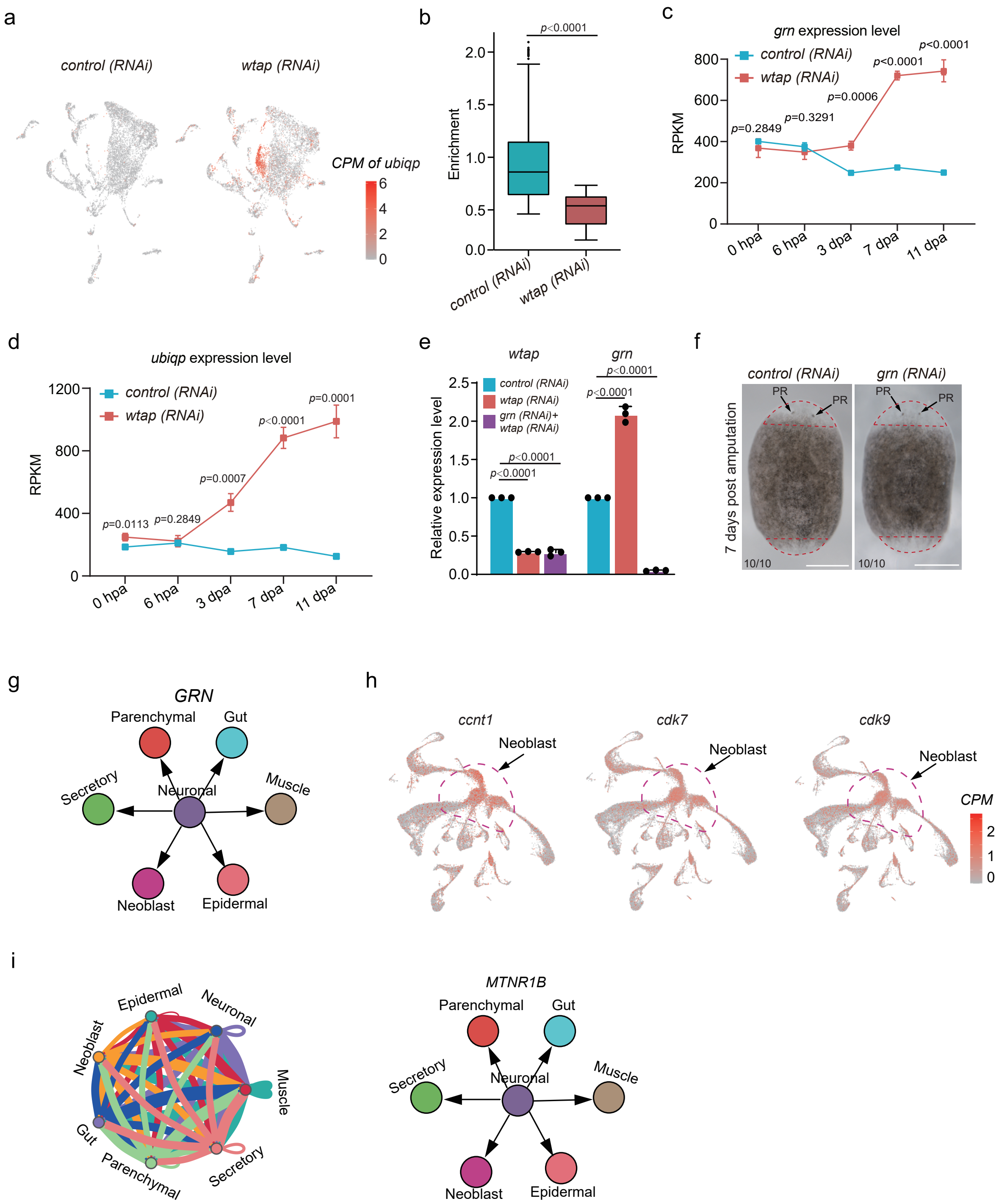

a
control (RNAi)
wtap (RNAi)
CPM of ubiqp
b
p<0.0001
Enrichment
control (RNAi)
wtap (RNAi)
c
grn expression level
control (RNAi)
wtap (RNAi)
RPKM
p=0.2849
p=0.3291
p=0.0006
p<0.0001
p<0.0001
0 hpa
6 hpa
3 dpa
7 dpa
11 dpa
d
ubiqp expression level
control (RNAi)
wtap (RNAi)
RPKM
p=0.0113
p=0.2849
p=0.0007
p<0.0001
p=0.0001
e
wtap
grn
control (RNAi)
wtap (RNAi)
grn (RNAi)+ wtap (RNAi)
Relative expression level
p<0.0001
f
control (RNAi)
grn (RNAi)
7 days post amputation
PR
10/10
g
GRN
Parenchymal
Gut
Secretory
Neuronal
Muscle
Neoblast
Epidermal
h
ccnt1
cdk7
cdk9
Neoblast
CPM
i
Epidermal
Neuronal
Neoblast
Muscle
Gut
Parenchymal
Secretory
MTNR1B

**Extended Data Fig. 5 | Identification of NP-like cell markers, Related to Figure 6.**

**a**, Expression level of *ubiqp* in *wtap* depleted (*wtap* RNAi) and normal (*control*) planarian during regeneration. **b**, The enrichment of $m^6A$ modification on *grn*. Data were analyzed by the two-tailed unpaired Student *t*-test. **c**, Lineplot showing the dynamic expression level of *grn* expression levels in both *control* (top) and *wtap* knockdown (bottom) planarians during 5 regeneration stages. The expression level is quantified from bulk RNA-seq. **d**, Expression level of *ubiqp* in *wtap* depleted (*wtap* RNAi) and normal (*control*) planarian during regeneration. Data were analyzed by the two-tailed unpaired Student *t*-test. **e**, qPCR showing the relative expression of *wtap* and *grn* in control (RNAi), *wtap* (RNAi) and double knockdown of *wtap* and *grn* at 7 dpa. **f**, Phenotype of normal (control) and *grn* depleted (*grn* RNAi) planarian at 7 dpa. Bottom left number, animals with phenotype of total tested. Scale bar, 500 µm. **g**, Network visualization showing NP-like cell communicates with other cell types through secreting GRN in *wtap* knockdown (RNAi) samples. **h**, Expression level of *ccnt1, cdk7* and *cdk9* in *wtap* knockdown (RNAi) samples. **i**, Network visualization of ligand-receptor pair numbers among different cell types (left). Network visualization of specific pairs among different cell types in *wtap* (RNAi) samples (right).
